# Single RNA molecule resolution spatial imaging of immunotherapy response in triple negative breast tumors harboring tertiary lymphoid structures

**DOI:** 10.1101/2024.03.19.585798

**Authors:** Richard H. Mebane, Teia Noel, Nathan Ing, Kenneth Gouin, Aagam Shah, David Zitser, Andrew Martinez, Gaorav Gupta, Alice Ho, Heather McArthur, Stephen S. Shiao, Simon. R.V. Knott

## Abstract

Cancer immunotherapy trials have had variable success, prompting a search for biomarkers of response. Tertiary lymphoid structures (TLS) have emerged as prognostic for multiple tumor types. These ectopic immunological bodies resemble secondary lymphoid organs with segregated B and T cell zones, but they are heterogeneous in their organization and cellular composition. These features have consequences in terms of prognostication and disease clearance, so there is interest in what drives TLS heterogeneity and corresponding immunological responses. We applied single RNA molecule resolution imaging to study biopsies from triple negative breast tumors harboring TLS where the biopsies were taken longitudinally, prior to therapy, after pembrolizumab and after pembrolizumab with radiation therapy. We developed a computational framework to identify TLS and tumor beds and to align spatial trajectories between the immune and malignant structures for systematic analyses. We identified two tumor types based on immune infiltration profiles in the tumor bed. Immune “infiltrated” tumors were eliminated after pembrolizumab, while “non-infiltrated” tumors saw gains in effector T cells and myeloid cells after pembrolizumab and were cleared after pembrolizumab with RT. TLS from infiltrated tumors had better separation of B and T cell zones and had higher expression of immunoreactivity gene pathways in most cell types. Further, malignant cell MHC expression was higher in the tumor beds of infiltrated tumors, providing one plausible mechanism for the groupings. In non-infiltrated tumors, classical dendritic cells enter the tumor bed from TLS proximal zones after pembrolizumab and bring transcription of the CXCL9 chemokine, which can recruit T cells and promote T cell effector phenotypes and was higher in infiltrated tumors at baseline.

## Introduction

The presence of tumor-infiltrating lymphocytes (TILs) within the tumor microenvironment (TME) is increasingly recognized as a favorable prognostic factor in various cancer types. These cells, when organized into tertiary lymphoid structures (TLS) with their distinct arrangement of B and T cells, are pivotal in predicting responses to anti-cancer treatments. TLS share anatomical similarities with secondary lymphoid organs like the spleen and lymph nodes, enhancing adaptive immune responses at their formation sites^1, 2^. The prognostic impact of TLS varies, with their location and maturity, as measured by segregation of B and T cell zones, influencing outcomes^3,4, 5^. Mature TLS, and those with high numbers of LAMP^+^ dendritic cells, MECA-79^+^ endothelial cells, and follicular helper T cells correlate with better disease-free survival and chemotherapy response^3, 6, 7, 8, 9, 10, 11, 12, 13^. Conversely, TLS containing immunosuppressive T regulatory (Treg) and B regulatory (Breg) cells are associated with poorer clinical outcomes^3, 14, 15, 16, 17, 18^. This variability underscores the critical need to understand programs guiding the architecture of TLS and how they change after different treatment regimens, as they hold profound implications for cancer prognosis.

Gene signatures from bulk RNA sequencing and single cell RNA sequencing studies have been used to examine immune cell programs within TLS, yet these technologies lack the spatial information required to fully elucidate the TLS cellular ecosystem^19, 20, 21, 22, 23^. Spatial transcriptomics has also been applied to study TLS but not in the form that allows single cell resolution measurements, and this was not done on paired pre- and post-treatment samples^24^. Several tissue staining protocols have also been applied to study broad cell type distributions in TLS from clinical trial samples, but these techniques did not allow cell type specific phenotypic analyses and did not shed light on the cellular phenotypes that exist within TLS or the phenotypic shifts that take place after treatment^20, 21, 22^.

To overcome the above-mentioned challenges, we applied single RNA molecule resolution imaging (CosMx) of 1000 gene targets to TLS harboring triple negative breast cancers (TNBCs) and developed a computational framework that integrates cell type annotation, niche calling and the identification of TLS, tumor bed and corresponding intermediate zones to examine both the structural and phenotypic differences that exist within each tumoral region. Importantly, the samples profiled encompass longitudinal biopsies (pre-treatment, on-treatment), where patients were treated with pembrolizumab or pembrolizumab in combination with radiation, so that we were also able to capture and describe treatment induced spatial and phenotypic shifts within the TME resulting from treatment with immunotherapy. This comprehensive framework aims to not only elucidate the dynamic interplay between various cell types within the TME and TLS, but also to provide insights into the mechanisms through which treatments, particularly immunotherapy and its combination with radiation, influence these cellular and molecular landscapes.

## Results

### Single molecule resolution imaging of TNBC harboring TLS

To examine how single RNA molecule imaging can delineate the details of the TME compared to other methods, we analyzed longitudinal biopsies from four TLS harboring tumors using the Nanostring CosMx platform and a panel of ∼1000 genes covering most genes identified as prognostic for immunotherapy trials previously^25^. The samples were obtained from a prior study that used scRNAseq and Co-Detection by Indexing (CODEX) on TNBC biopsies taken before therapy, after pembrolizumab (pembro) and after pembro + RT (Fig S1A), that identified a responsive patient population harboring tumors constituted by CD20^+^ B cells that formed localized clusters spatially correlated with T cells, thereby resembling TLS^25^. On average, this group showed T cell clonal expansion prior to therapy and epithelial cell elimination after pembro. After quality control filtering for cells with sufficient molecules per cell, and genes exceeding the median negative probe rate per cell, 2.78 x 10^8^ RNA molecules were analyzed across 651,683 single cells, with an average of 427 molecules and 193 genes being measured in each cell (Methods).

To annotate the imaged cells, we transferred cell type labels from corresponding scRNAseq data using the scANVI software package (Methods, Fig S1B-D)^25, 26^. Spatial correlations were high amongst B cells, T cells as well as amongst both classical and plasmacytoid dendritic cells (cDC and pDC). While normal epithelial cells (epi. n.) also showed some spatial co-localization with these immune subsets, malignant epithelial cells (identified previously based on inferred copy number changes, epi. m.), appeared to occupy a distinct tumor zone (Fig 1B)^25,27^. To formalize this observation, we created cellular niches in which we calculated broad cell type proportions within an average radius of 2 cells of each cell and then leiden clustered the resultant profiles (Fig 1C, Methods)^28^. Niches 0, 3 and 6 were dominated by myofibroblasts (myCAFs), with niche 0 also showing macrophage presence and with niche 6 showing inflammatory fibroblast (iCAF) inclusion. Niche 1 was constituted mainly by epi. m. cells, while niche 5 was most enriched for epi. n. Niche 2 contained many endothelial cells, while niches 4 and 7 harbored mostly immune cells. Niche 4 was constituted mainly by T and B cells, while niche 7 contained more plasma cells. Beyond the fibroblast rich niches 0, 3 and 6 the identified cellular communities did not show strong inter-niche spatial correlation patterns (Fig 1D).

**Figure 1.**
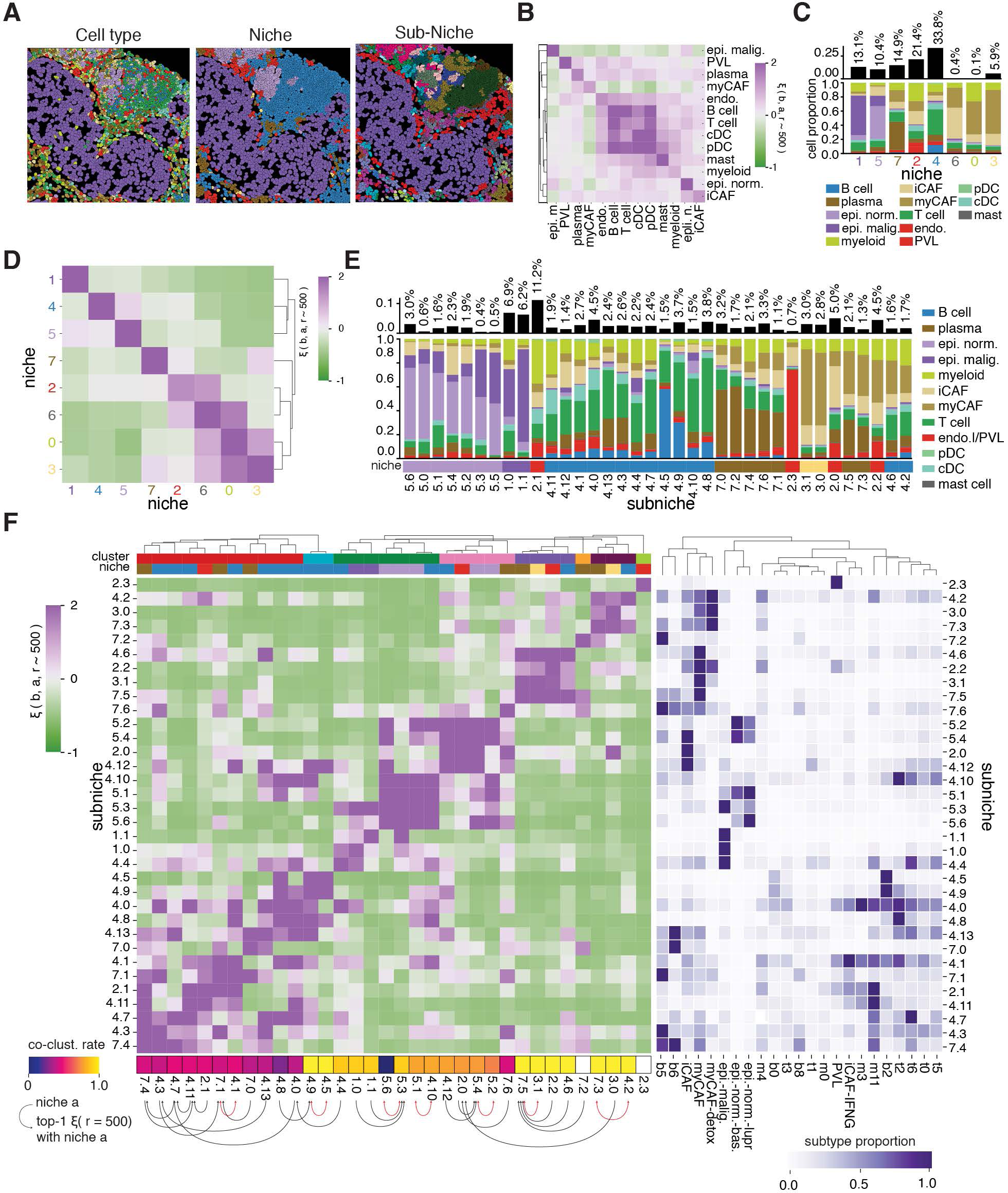
**A.** Example cell segmentation images colored by cell type, broad niche, and sub-niche. **B.** Spatial cross-correlation coefficient for each cell type pair at a fixed scale of 500 coordinate units. Positive values correspond to an overdensity relative to the background. **C.** Bar plot of the percentage of cells assigned to each broad niche (top); Stacked bar plot of the percentage of cells in each niche belonging to each cell type class (bottom) **D.** Spatial cross-correlation coefficient for each broad niche pair at 500 coordinate units. Niches are highly clustered with themselves at this scale. **E.** Bar plot of the percentage of cells assigned to each subniche (top); Stacked bar plot of the percentage of cells in each subniche belonging to each cell type class (bottom). **F.** Spatial cross-correlation coefficient for each sub-niche pair at a scale of 500 coordinate units along with subtype proportions of each sub-niche. Spatial sub-niche clusters are annotated at the top. Co-clustering rate at the bottom was determined by computing these spatial cross-correlations on 10,000 samples bootstrapped at the scale of individual fovs and averaging the rate at which each sub-niche was in the same spatial cluster as each other sub-niche (R); Heatmap of Min-max scaled cell subtype proportion in each subniche (L).

Single RNA molecule resolution imaging also allows for deeper cellular subclassifications, so we also applied the scANVI algorithm to label cells by subtype (Methods) and identified corresponding “subniches” using these labels and similar methodologies to those outlined above (Methods, Fig 1A, 1E, S1E & S1F)^26^.

Interestingly, even when viewed through the lens of their broad cell constituents, spatial transcriptomics allowed us to identify multiple subniches generated from the same niche. For example, the subniches that were derived from broad niche 4 segregated based on their ratio of T to B cells, with subniches 4.5 and 4.9 showing the highest abundance of the latter cell type (Fig 1E). Additionally, subniches 4.2 and 4.6 had high myCAF and iCAF involvement (Fig 1E). Another notable distinction was niche 2, which segregated into subniche 2.1, 2.2 and 2.3, where 2.1 contained substantial macrophage numbers, 2.2 contained high fibroblast counts and 2.3 contained mostly endothelial cells.

Subniches allowed us to identify spatial relationships between communities that were distinct from the relationships emanating from the same broad niche. For example, one large spatially defined cluster of subniches was identified that contained most of the T and plasma cell rich subniches derived from niche 4 as well as subniche 7.0 and 7.4 from niche 7. In contrast, iCAF subniche (4.12) and naïve T cell subniche (4.10) were found co-localized with niche 5 subniches containing mainly epi. n. Notably, the two B cell subniches (4.5 and 4.9) were found most spatially correlated with each other, and both also showed strong spatial association with naïve T cell subniche 4.8. (Fig 1F)

### Determining how architectural context influences cell type specific gene expression programs

To study how architectural context influences gene expression programs, we developed a methodology using a systematic pairwise analysis of subniches, for each pair analyzing gene expression differences in each cell type. We measured the distinctiveness between subniche expression programs adapting Augur AUC scores (Methods), and then correlated these AUC scores with the corresponding spatial correlation values for the subniche pairs (Fig 2A)^29^. This found there was a higher discriminatory capacity for cells when they occupied subniche pairs that were not spatially correlated. This anticorrelation between AUC scores and spatial relatedness was observed for all cell types except for endothelial cells and cDCs, potentially indicating these later two cell types are not susceptible to changing in response to ligand cues in the tumor microenvironment.

**Figure 2.**
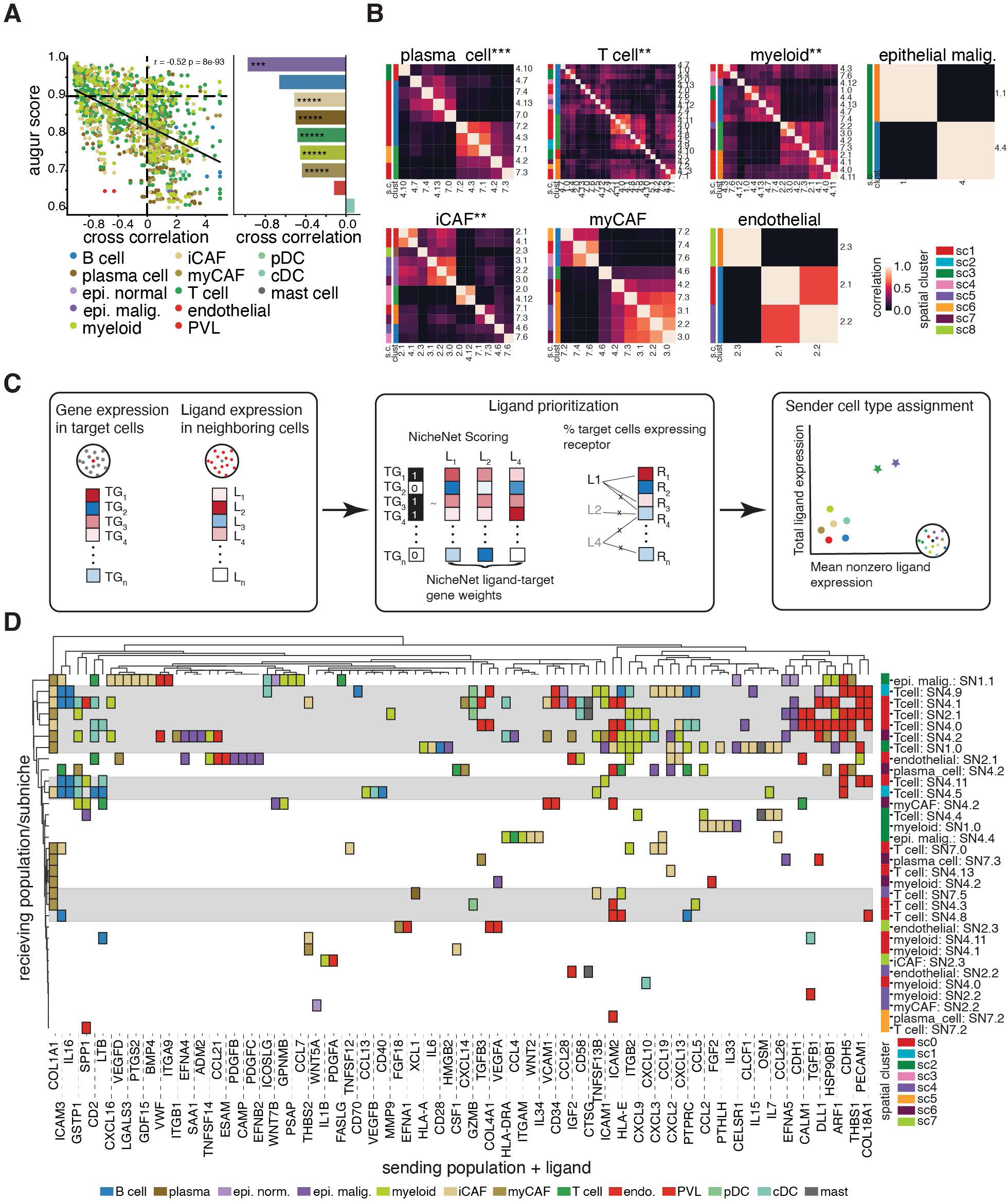
**A.** Scatter plot of spatial cross-correlation coefficient and gene expression augur score for each sub-niche pair of a given broad cell type. Each dot represents a cell type/subniche pair, colored by cell type. Pearson correlation coefficient and p-value for testing non-correlation are reported above (L); Bar plot of Pearson correlation coefficients of the relationships shown in the scatter plot, per cell type. Bars are colored by cell type and asterisks indicate the level of significance of Pearson correlation (* <= 0.1, ** <= 0.05, *** <= 0.01, **** <= 10^-3^, ***** <= 10^-4^) (R). **B.** Heatmaps of jaccard scores comparing gene signatures describing each cell type/subniche context. Rows are annotated by broad niche, subniche spatial clusters reported in Fig 1F, and clustering of jaccard scores. Asterisks indicate the level of significance resulting from a Chi-squared-contingency test, testing the overlap of jaccard score clusters with spatial clusters (* <= 0.1, ** <= 0.05, *** <= 0.01, **** <= 10^-3^, ***** <= 10^-4^). **C.** Schematic of ligand-receptor analysis used throughout this study. First, we identify a gene expression program in some target cells through differential expression, spatial patterns, etc. Then, we find ligands expressed in cells neighboring these target cells. Finally, target genes are compared to NicheNet predictions for each ligand to determine which ligand in which cell type is likely influencing target expression. **D.** Ligands explaining gene expression programs found to differentiate cell types in different subniches. Colors indicate the most-likely sending cell type population of each ligand (columns) affecting the downstream genes of each cell type-in-subniche target population (rows). Rows are color-annotated by spatial clusters described in Fig 1F.

We then compared subniche specific gene expression programs and their relation to the spatial clusters identified above. For each cell type we calculated pairwise jaccard similarity scores between pairs of subniches. These scores were then clustered to identify groups of subniches that showed high similarity in their defining gene expression programs for specific cell types (Fig 2B). Application of the chi-squared contingency statistic, to test if resultant gene program clusters were related to the spatial clusters, found that spatial clusters were predictive of gene program clusters for multiple cell types including plasma cells, T cells, macrophages and iCAFs (Fig S2B).

We next wanted to explore whether there were ligand-receptor interactions associated with the cell states identified for each spatial cluster. For each cell type, we used the trained Augur models from above to identify cell types that differed phenotypically between subniches (AUC > 0.9) and treated these as receiving cell populations for downstream analysis. For each receiving population we then identified receptors listed in the Nichenet database that were expressed in more than 20% of cells within one of the subniche pairs and where the corresponding ligands were overexpressed by neighboring cells in the same subniche, regardless of cell type^30^. Identified ligands were then scored on the receiving cell gene signature using the NicheNet database (Methods). Finally, for the ligands showing potential to drive the receiving cell state, we applied isolation forests to determine the most likely “sending” cell population (Fig 2D, Methods)^31^. In our samples, we found that many of the subniches harbored within spatial clusters 0, 1 and 2 were involved in similar ligand-receptor guided communication patterns. Notably, T cells from these zones appeared to be prominently guided in their expression programs by endothelial cells through ligands such as ARF1, and PECAM1, as well as through CAF and myeloid expressed CXCL chemokines, CAF expressed COL1A1 and ICAM2 as well as IL16 expressed by B cells (Fig 2D, Supplementary Data 1).

### Identification and transcriptional characterization of TLS and associated tumoral zones

In addition to our unbiased niche-based analyses, we also directly looked for and analyzed tumor regions corresponding to TLS and the tumor beds these structures are intended to eliminate. Besides endothelial cells, B cells showed the most distinctive spatial distribution, with their maximal enrichment being in subniches 4.5 and 4.9 (Fig. 1E). Spatial correlation analyses also found these two subniches were reciprocal nearest neighbors, and that they were frequently abutted by naïve T cell enriched subniche 4.8 (Fig. 1F). Based on these findings we reasoned 4.5 and 4.9 likely represented the TLS “B cell zone” (BCZ) and that subniche 4.8, when next to these zones represented the “T cell zone” (TCZ) of TLS (Fig. 3A). We also found co-located malignant cell enriched subniches 1.0 and 1.1 together encompassed (∼74%) of baseline malignant cells and reasoned that these zones represent malignant cell zones (MCZ) therapeutic interventions were intended to eliminate (Fig. 1E-F, 3A).

**Figure 3.**
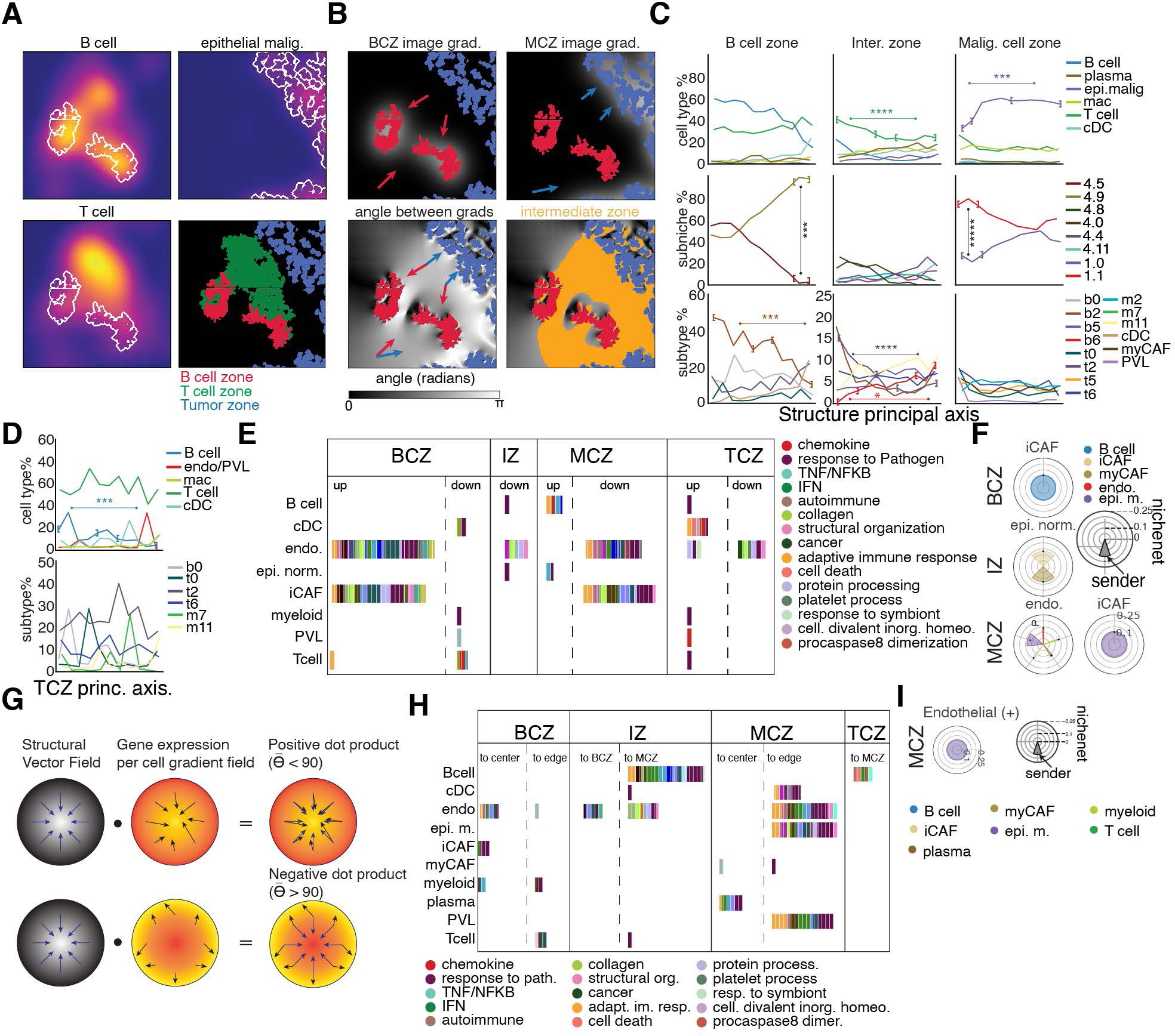
**A.** Images of B cell, T cell, and epithelial tumor zones. Background images show the density of these cells in the same region of tissue, and the white outlines show the location of each connected cluster of cells. **B.** Example of the determination of the intermediate zone bridging the B cell TLS core and the malignant cell zone. First, we compute the gradient fields of the convolved images of each zone. Then, we calculate the angle between the gradients at each pixel, setting a minimum cutoff of 0.8π radians for the intermediate region. **C.** Proportions of broad cell type (top), sub-niche (middle), and cell subtype (bottom) moving through each zone from the core of the B cell zone on the left, through the intermediate zone in the center, and ending at the core of the malignant cell zone at the right. We include only those trajectories which account for at least 10% of any one spatial bin. Asterisks indicate statistical significance of a Wilcoxon rank sum test between proportions in the bracketed regions (* <= 0.1, ** <= 0.05, *** <= 0.01, **** <= 10^-3^, ***** <= 10^-4^). **D.** Proportions of broad cell type (top) and subtype (bottom) moving through TLS T cell zones connected to B cell zones. Asterisks indicate statistical significance of a Wilcoxon rank sum test between B cell proportions in the bracketed region (* <= 0.1, ** <= 0.05, *** <= 0.01, **** <= 10^-3^, ***** <= 10^-4^) **E.** Pathways associated with gene expression programs in each cell type in each zone found through differential expression between each cell type in a given zone and the rest. (GSEA; FDR < 0.05). **F.** Radar plots describing ligand-receptor analysis results for this differential expression. Each plot corresponds to a cell type/zone target population. Color of bars represents cell type sender, radius represents maximal nichenet score across all ligands sent by cell type, dots represent median nichenet score across all ligands sent by celltype, and width of bar represents percentage of all interactions coming from cell type. Interactions included here are those with a NicheNet score > .1. **G.** Example of gradient analysis used throughout this work. We first compute the gradient of the convolved image of one instance of a particular zone. Then, we compute the gradient of expression of each gene in each cell type in this region and divide it by the local density of that cell type. Finally, we take the dot product between these two vector fields in order to find the angle between them, identifying genes which increase or decrease in expression per cell as you move toward the center of an instance. **H.** Pathways associated with gene expression programs found to move together through the previously described gradient analysis. (GSEA; FDR < 0.05). **I.** Radar plots illustrating ligand-receptor analysis results which describe ligands in sending populations that affect groups of genes with similar gradients in receiving populations. Interactions shown here are those with a NicheNet score > .1.

To formally define intermediate zones (IZ) residing between BCZ and MCZ, we performed gaussian image convolutions of full biopsies based on BCZ and MCZ coverage, separately, and then computed the gradient vector field of the resulting convolved images (Fig. 3B). We then computed the angle between the BCZ and MCZ gradient fields in each biopsy at per-pixel resolution and set a cutoff angle of 0.8π to define pixel locations constituting each BCZ-MCZ combination, and also setting a 400-coordinate unit distance cutoff to ensure intermediate zones only connect nearby BCZs and MCZs (Fig. 3B).

To examine how the TLS associated structures interact, we used the gradient fields originally employed to define IZs to define trajectories moving from the center to the periphery of BCZs and MCZs and for IZs from the edge of the BCZ to the edge of the corresponding MCZ. Then, to study cellular proportions and how they changed across regions, for each BCZ, IZ and MCZ, we defined 10 equally spaced bins across the corresponding gradients and calculated broad cell type, cell subtype and subniche proportions per bin. We found memory B cells (b2) showed the strongest spatial pattern within BCZ, going from heavily to weakly enriched from the core to the periphery (Fig 3C). This also corresponded to a gain in subniche 4.9 dominance near the periphery. The TCZ saw no change, outside of a modest drop in B cell proportions moving away from the BCZ (Fig 3D). Looking further out from the BCZ along the IZ gradient, T cells did drop in proportion moving towards the MCZ, and this coincided with a drop in naïve T cells (t2, Fig 3C). Along the same trajectory there were gains in IGHG1 expressing plasma cells (b6). Finally, moving from the MCZ edge to the inner core there was a rise in epi. m. and subniche 1.0 proportions coincident with a drop in subniche 1.1, the more heterogeneous of the two malignant cell zones (Fig 3C).

We next performed cell type specific differential gene expression comparisions for each zone, and applied gene set enrichment analysis (GSEA) to understand the functional phenotype of individual cells within these intratumoral structures (Methods). Compared to the other regions, BCZ endothelial cells, iCAFs and T cells all showed upregulation of gene modules related to adaptive immune response, and these pathways were down in the former two cell types in MCZs (Fig. 3E). Other gene sets showing the same trends in endothelial and iCAFs were related to MHC, protein processing, response to pathogens and metabolism. Additionally, we associated genes enriched in each cell type and spatial zone with ligand-receptor interactions. This determined BCZ iCAFs were potentially being influenced by B cells, while in the MCZ iCAFs and endothelial cells appeared to be regulated by interactions with epi. M (Methods, Fig 3F, S3, Supplementary Data 2 & 3).

### Intrastructural spatial expression patterns reflect interstructure relationships

To study spatially associated gene expression differences across TLS associated structures, we performed a convolution for each gene and cell type combination in each image, divided the image pixels by the local density of that cell type and then computed a corresponding gradient (Fig 3G). The resulting vector field represents the direction of positive change of expression of one gene in a single cell type per cell. Finally, for each pixel in the image we computed the component of the gene expression gradient along the direction of the principal axis of the given structure that was computed previously.

We then performed a Kolmogorov-Smirnov test comparing the resulting distribution with a uniform distribution to identify genes that increase or decrease in the direction of the principal axis of any structure type. This found adaptive immune response and response to pathogen signatures in BCZ endothelial cells and iCAFs showed increases moving towards the BCZ core, along with an interferon gene signature (Fig 3H). Following the same gradient, myeloid cells saw an increase in genes related to a general and innate immune response, while T cells showed increases in complement and metabolism genes moving towards the BCZ periphery.

In the intermediate zone, B cells showed expression-based increases in adaptive immune response, response to pathogen, TNF/NFKB signaling, interferon signaling, metabolism and MHC expression moving from the BCZ towards the MCZ (Fig 3H). Finally, a variety of cell types showed spatially defined changes within the MCZ. For example, cDCs, endothelial cells, epi. m. and PVLs all showed increased expression of signatures related to adaptive immune response, MHC and response to pathogen moving towards the MCZ periphery, whereas plasma cells showed changes in the same pathways but moving in the opposite direction.

We next examined whether cellular interactions may be influencing intrazone expression trajectories. To address this, for each cell type in each zone we defined positive gradient signatures as genes passing the 95th percentile of positive dot products with respect to the zone principal axis, and negative gene gradients as those in the bottom 5th percentile of the negative dot products. For each pair of cell types, we looked for an association of ligands in one cell type and genes in the second with the same directionality. We filtered for ligands associated with receptors expressed in at least 30% of the target cell population. Lastly, we ensured that ligand expression in the neighborhoods of the target cell population correlated with the signature scores of the target cells. This found several of the interzone relationships described in Fig 3F changed spatially within zones. For example, B cell influence on endothelial cell programs increased moving to the periphery of the BCZ, like what was also seen with epi. m. influence on endothelial cells and iCAFs in MCZs (Fig 3I). Indeed, most intra-zone spatially defined interactions increased in strength moving towards the periphery of the corresponding zone (Supplementary Data 4 & 5).

### TLS harboring tumors display heterogeneity in their response trajectories

The four tumors chosen for single RNA molecule resolution imaging analysis were selected because they represent two distinct response trajectories in terms of epi.m. elimination, despite all four being TLS harboring tumors. Patients pt02 and pt16 had Niche 1 elimination after pembro alone, but in pt12 and pt43 tumor cell elimination was not seen until after the combination therapy (Fig 4A). To study if these tumors could be distinguished based on their molecular characteristics, we clustered zones of the same type by their constituent cellular proportions. MCZ instances separated by the response trajectories of their corresponding tumor, mainly due to more T cells and myeloid cells being present in MCZ instances from pt02 and pt16 (Fig 4B). Due to this, for brevity moving forward we call tumors from pt02 and pt12 “infiltrated” and lesions from pt16 and pt43 “non-infiltrated”, though other cell types like endothelial cells, PVL and cDC were enriched in tumors from the latter group. Infiltrated and non-infiltrated samples also segregated by BCZ cellular proportions (Fig 4B). Infiltrated BCZs were mainly composed of B and T cells, while non-infiltrated tumors had higher proportions of cDC, pDC and myeloid cells amongst other types. A weaker segregation by infiltration status was observed when IZ and TCZ were examined, but differences were identified, including increased T cell proportions in non-infiltrated IZs and higher epi. m. myeloid and plasma cell proportions in infiltrated tumors. TCZs from infiltrative tumors had more T cells, while non-infiltrated tumors had more pDC and endothelial cells.

**Figure 4.**
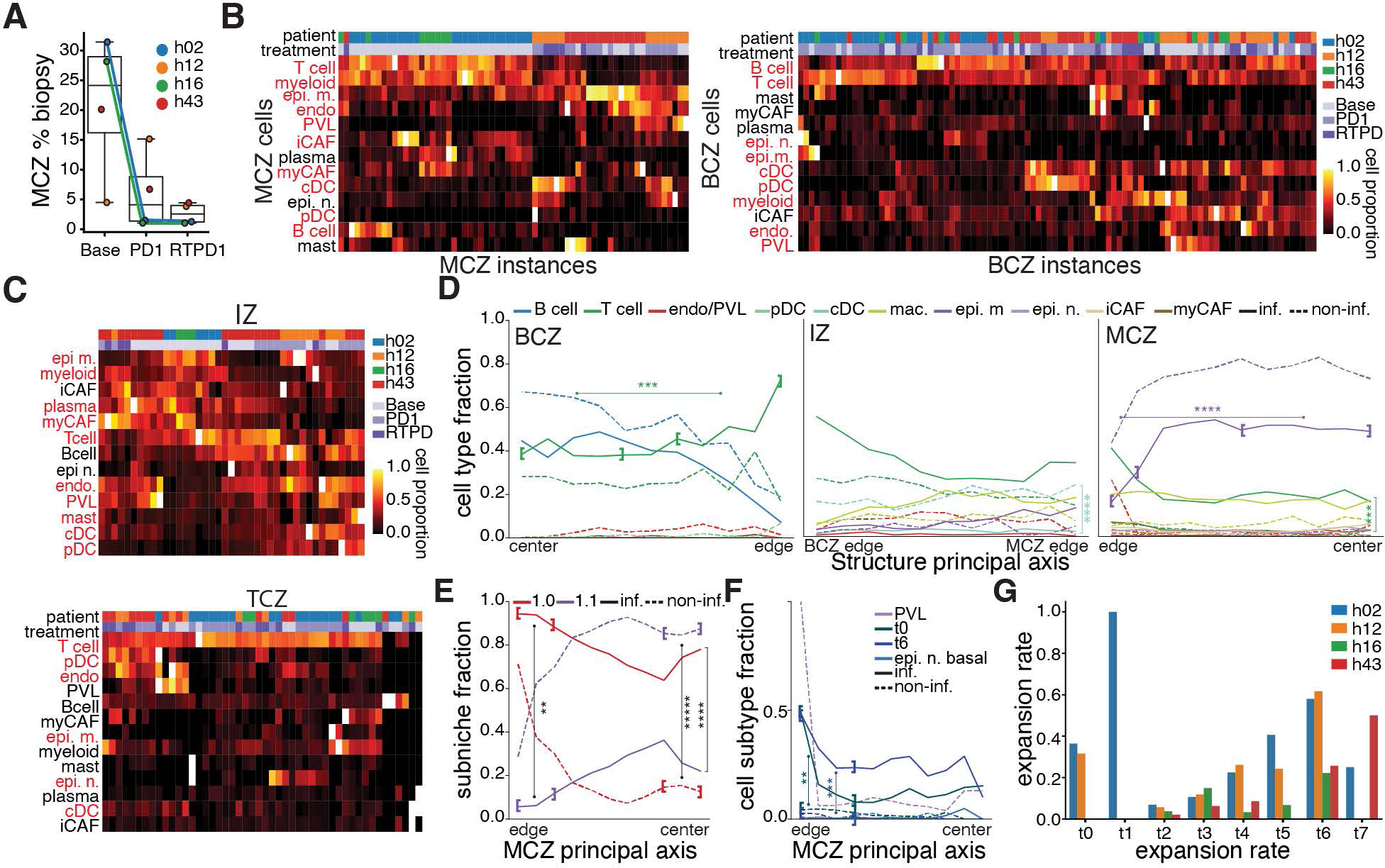
**A.** Box plots of percentage of total cells in each biopsy consisting of niche 1 cells, split by time point. Points are colored by patient. **B.** Heatmaps showing broad cell type compositions of each instance of malignant cell zones (L) and B cell zones (R). Columns are color-coded by patient and treatment. Cell types highlighted in red indicate where we find a statistically significant difference in infiltrated versus non-infiltrated instances (Mann-Whitney U test; p <= .05). **C.** Heatmaps of cell type compositions of each instance of the intermediate zone and T cell zone. **D.** Cell type proportions traveling through each zone for infiltrated (solid) and non-infiltrated (dashed) patients at baseline. Trajectories shown are those with a statistically significant difference between infiltrated and non-infiltrated patients in at least two consecutive spatial bins. (Mann-Whitney U test; p <= .05). Asterisks indicate statistical significance of a Wilcoxon rank sum test between proportions in the bracketed regions (* <= 0.1, ** <= 0.05, *** <= 0.01, **** <= 10^-3^, ***** <= 10^-4^). **E.** Sub-niche proportions traveling through the malignant cell zone at baseline for patients of each infiltration status. Sub-niches shown must pass the same test as the previous plot. Asterisks indicate statistical significance of a Wilcoxon rank sum test between proportions in the bracketed regions or the entire region where a bracket is not indicated (* <= 0.1, ** <= 0.05, *** <= 0.01, **** <= 10^-3^, ***** <= 10^-4^). **F.** Cell subtype proportions through the malignant cell zone. Subtypes shown pass the same test as the previous plot. Asterisks indicate statistical significance of a Wilcoxon rank sum test between proportions in the bracketed regions (* <= 0.1, ** <= 0.05, *** <= 0.01, **** <= 10^-3^, ***** <= 10^-4^). **G.** T cell expansion rates from the single cell data of each patient in this study at baseline.

Most TLS-structure associated proportional differences between infiltrated and non-infiltrated tumors were uniformly distributed along the principal axis of the corresponding structures (Fig 4D). However, examining B and T cell rates near the BCZ edge and early on within IZ, it appears infiltrated tumors have more mature TLS with stronger segregation of T and B cell zones.

Within the MCZ, the highest cellular heterogeneity was at the periphery for both infiltrated and non-infiltrated tumors. Both tumors also saw epi. m. proportions become the dominant population moving towards the core, but this was less obvious in the latter tumor type (Fig 4D). Heterogeneity at the MCZ periphery was matched with higher subniche 1.0 proportions in both tumor groups but this appeared driven by T cell infiltration only in infiltrated tumors (Fig 4D, 4E). Furthermore, subniche 1.0 dominates all along the principal axis of infiltrated MCZ, but 1.1 becomes the most prominent subniche in non-infiltrated tumors early when traversing along this trajectory. In terms of subtypes, the only significant differences between infiltrated and non-infiltrated tumors were in the MCZ, where effector T cell subgroups t0 and t6 were found near the edge of infiltrated tumors (Fig 4F). Importantly, when examining the corresponding scRNAseq and paired TCR data we found infiltrated tumors had higher expansion rates in almost all T cell subgroups at baseline, but this was particularly noticeable in t0 and t6, as well as the follicular helper CD4^+^ T cell population t4 (Fig 4G).

### Transcriptional differences between infiltrated and non-infiltrated tumors

Infiltrated MCZ house endothelial cells, epi. n, epi. m, myCAFs, macrophages and T cells with higher expression of genes related to adaptive immune response and response to pathogens, while cDC, iCAF, myCAF and macrophages in non-infiltrated MCZs are enriched for genes involved in structural organization (Fig 5A). Additionally, iCAFs in this zone as well as in the BCZ and IZ of non-infiltrated tumors are enriched for genes related to collagen and protein processing. Finally, another notable finding was infiltrated tumor T cells in all compartments display a chemokine related gene signature, with the most significantly upregulated gene driving this pathway being CXCL13 in all regions but BCZ (Fig 5A & S4D-F).

**Figure 5.**
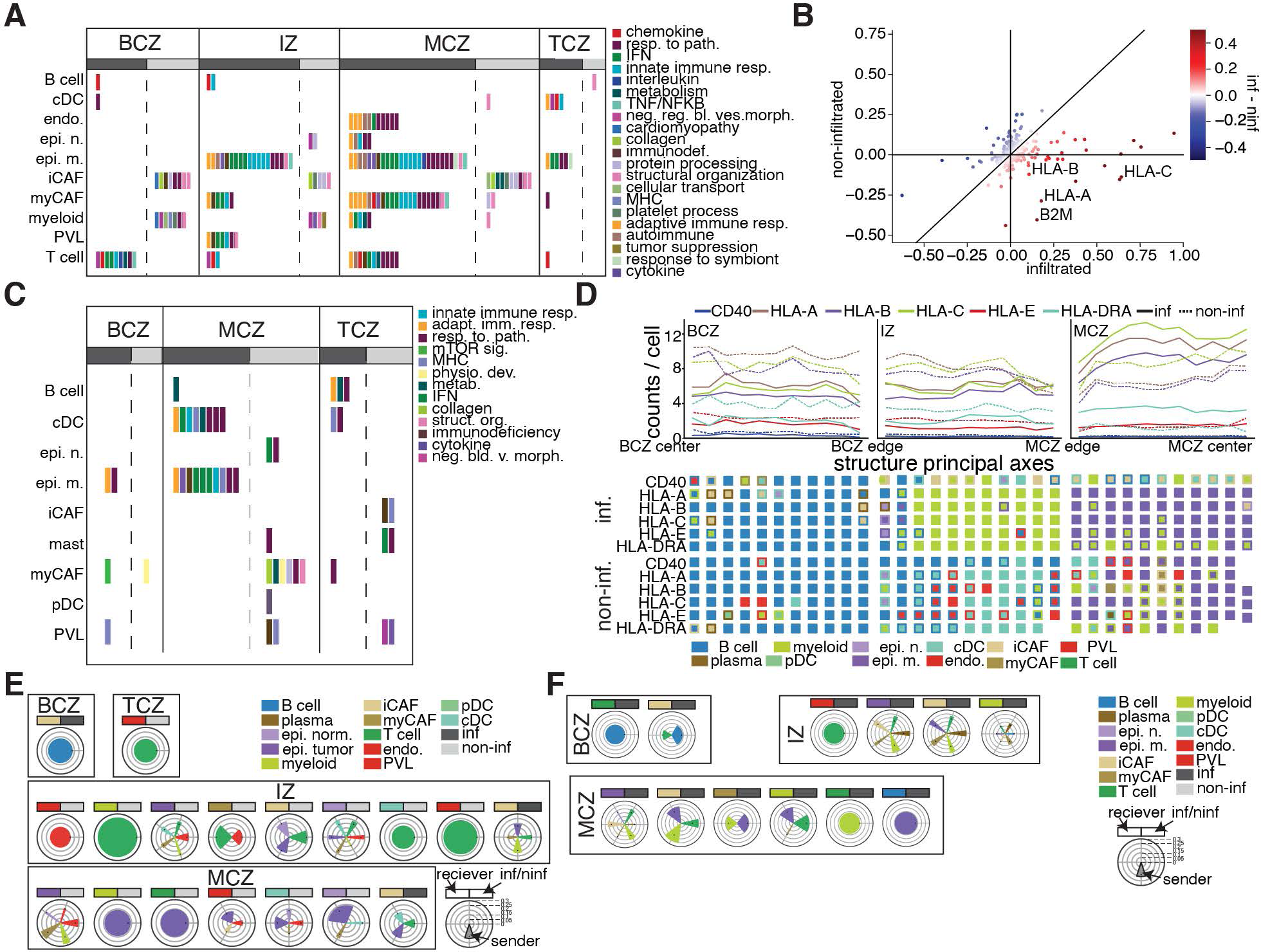
**A.** Gene expression programs found from differential expression between immune infiltrated and non-infiltrated biopsies at baseline in each zone. (GSEA; FDR < 0.05). **B.** Scatter plot of gradient differences between infiltrated and non-infiltrated patients. We compute the difference between the mean dot product of gene expression per cell in patients of each infiltration status and use this as the basis for our pathway analysis. **C.** Gene expression programs found from these gradient differences. (GSEA; FDR < 0.05). **D.** Counts per cell of immune checkpoint genes for infiltrated (solid) and non-infiltrated (dashed) patients through each zone. Heatmap below shows primary (outer) and secondary (inner) expressors of each gene in each spatial bin via the isolation forest method. [what determines which trajectories are shown] **E.** Ligand-receptor analysis for differential expression and **F.** gene gradient signatures. All results show interactions corresponding with a NicheNet score > .1.

We also studied how gene expression gradients within structures differed between infiltrated and non-infiltrated tumors. To do this, we computed the difference in their dot products with the zone principal axes and classified genes as differential if these dot products showed strong differences (Fig 5B & C). Though fewer related pathways were identified with this analysis, there was overlap with the findings reported in Fig 5A. For example, genes that show a greater upwards trend towards the MCZ center in epi. m. in infiltrated tumors were related to interferon signaling, innate immune response and response to pathogen (Fig 5C). One notable finding that could partially explain the presence of T cells within the infiltrated tumor MCZ at baseline was a difference in MHC expression patterns by epi. m, with many of the upregulated pathways for this cell type being driven by these genes (Fig 5A-C). Interestingly, when MHC molecule expression patterns were further examined, we found higher expression in IZ of non-infiltrated tumors, driven by B cells and cDC (Fig 5D & 4D).

B cells were the predicted ligand senders driving most of the general and intrastructural spatially defined differences measured in BCZ (Fig 5E, 5F, & S4A-C; Supplementary Data 6 & 7). In contrast, T cell expressed ligands were predicted to be the cause of most differences measured in the TCZ and IZ, while epi. m. ligands appeared to have the most influence on MCZ expression programs. This was particularly true in non-infiltrated tumors where epi m. were predicted to influence expression of macrophages and T cells found depleted in these zones through ligands such as ARF1, CXCL16, HSP90B1, IL34, VEGFB, THBS1 and CALM1.

### Pembrolizumab alters T cell phenotypes and infiltration status in TLS harboring tumors

In studying PD-L1 and PD-L2 expression, we found no overall expression difference between infiltrated and non-infiltrated tumors in BCZ, IZ or MCZ. However, the predicted senders of these signals were distinguishing (Fig 6A). In infiltrated BCZ and IZ, macrophages were predicted to be the primary cell type expressing PD-L1 and iCAFs were identified as the main senders of PD-L2. In contrast, cDCs were predicted to be the major expressors of both checkpoint ligands in non-infiltrating tumors. Additionally in infiltrated MCZ, malignant cells, macrophages and cDC are the predicted PD-L1 expressors and macrophages and iCAFs the predicted PD-L2 expressors, whereas in non-infiltrated lesions, epi. m. cells were the most frequently identified producers of both molecules.

**Figure 6.**
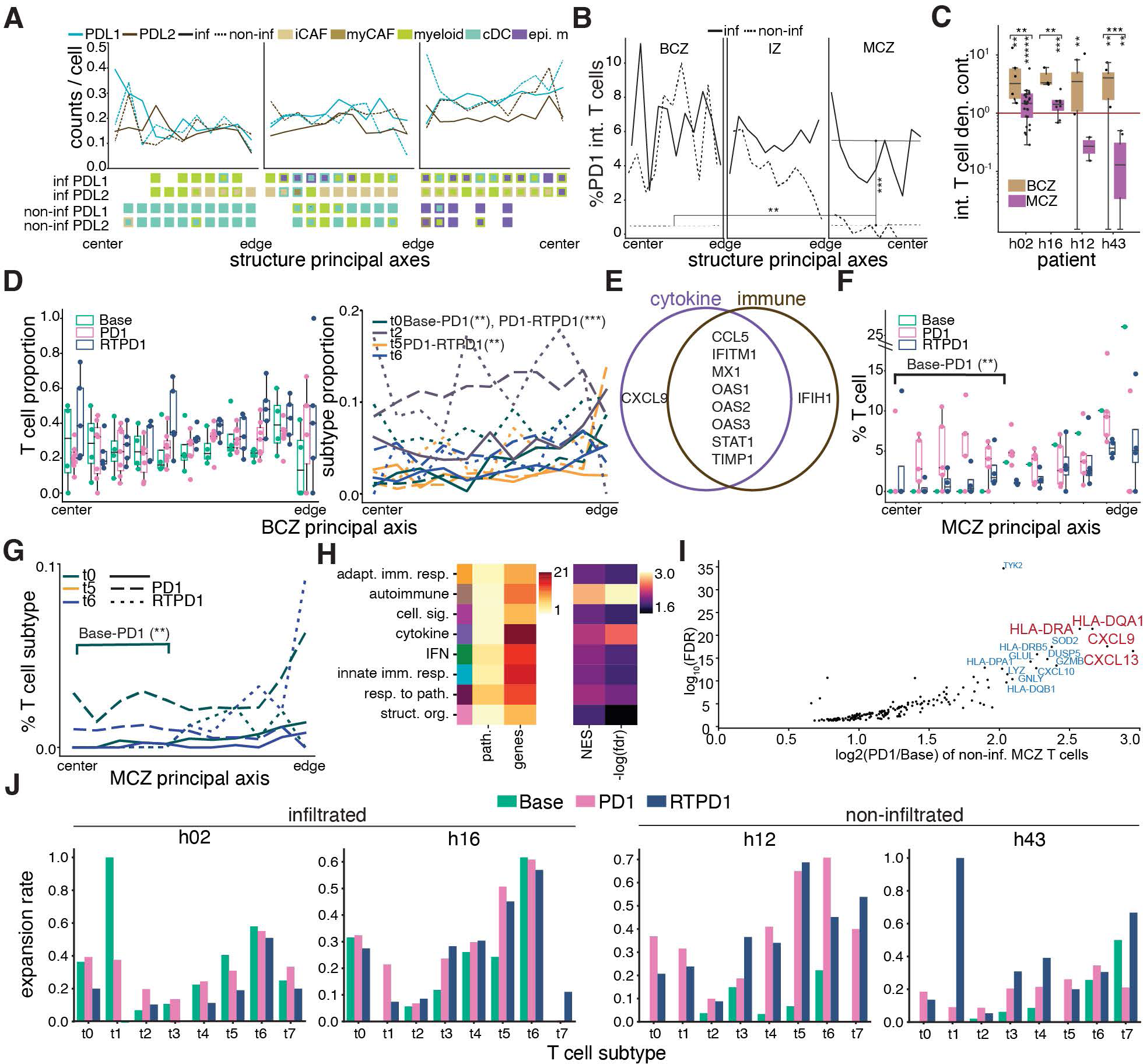
**A.** PDL1/2 counts per cell through each zone at baseline for infiltrated (solid) and non-infiltrated (dashed) patients. Isolation forest results for primary (outer) and secondary (inner) broad cell type expression in each spatial bin shown below. **B.** Proportion of PD1 interacting T cells moving through each zone. As you move through the intermediate zone into the malignant cell zone, PD1 interactions drop significantly in non-infiltrated patients. Asterisks indicate statistical significance of a Wilcoxon rank sum test between proportions in the indicated regions (* <= 0.1, ** <= 0.05, *** <= 0.01, **** <= 10^-3^, ***** <= 10^-4^). **C.** Box plots of density contrast of PD1 interacting T cells (T cells expressing PD1 and neighboring a cell expressing PDL1/2) and non-interacting T cells in the MCZ and BCZ instances at baseline, per patient. Asterisks indicated statistically significant differences between the BCZ and MCZ. **D.** T cell subtype proportions moving through the B cell zone for infiltrated and non-infiltrated patients. Trajectories shown have a statistically significant difference (p < 0.05) in at least two consecutive spatial bins between timepoints. Asterisks indicate statistical significance of a Wilcoxon rank sum test between proportions in full zone between the indicated time points (* <= 0.1, ** <= 0.05, *** <= 0.01, **** <= 10^-3^, ***** <= 10^-4^). **E.** Venn diagram of leading edge genes associated with cytokine and immune pathways for T cells in the malignant cell zone. (GSEA; FDR < .05). **F.** Box plots of overall T cell proportions moving through individual instances of the MCZ at each timepoint in non-infiltrated tumors. Asterisks indicate statistical significance of a Wilcoxon rank sum test between proportions in the bracketed region between the indicated time points (* <= 0.1, ** <= 0.05, *** <= 0.01, **** <= 10^-3^, ***** <= 10^-4^). **G.** T cell subtype proportions moving through the MCZ at each timepoint. Subtypes shown must have a statistically significant difference between at least two timepoints in at least two consecutive spatial bins. (Mann-Whitney U test; p <= .05). Asterisks indicate statistical significance of a Wilcoxon rank sum test between proportions in the bracketed region between the indicated time points (* <= 0.1, ** <= 0.05, *** <= 0.01, **** <= 10^-3^, ***** <= 10^-4^). **H.** Pathway summaries for T cells in the malignant cell zone at PD1 in non-infiltrated patients. Heatmaps show the number of pathway hits for each summary (GSEA; FDR < .05), number of unique genes in these pathways, GSEA normalized expression score, and FDR. **I.** Upregulated genes in malignant cell zone T cells at PD1 compared to baseline (LFC > .2, FDR < .05, and percent expressed in either group > .25) **(J)** T cell subtype expansion rates for each patient at each timepoint from the single cell data.

PD-L1 and PD-L2 expression gradients combined with T cell infiltration patterns resulted in the rates of PD1^+^ T cells interacting with PD-L1^+^ or PD-L2^+^ cells being similar across all zones in infiltrated tumors but dropping in non-infiltrated tumors moving from BCZ to MCZ cores (Fig 6B & S5A). Furthermore, there was a higher-than-expected rate of these interactions in BCZ relative to the rest of the tumor in both infiltrated and non-infiltrated samples, but this measure was lower than expected in non-infiltrated MCZ (Fig 6C).

In terms of T cell associated changes induced by therapy, BCZs from infiltrated tumors did not show T cell or T cell subtype proportional changes over the course of therapy. However, in non-infiltrated BCZ there were increased T cell proportions overall and increased t0 effector and t5 follicular helper subpopulations after addition of RT (Fig 6D & S5B-C). Modest gains were also seen in naive T cells (t2). No pathways were altered in BCZ T cells from infiltrated tumors after therapy, but cytokine and general immune related signatures were upregulated in BCZ T cells from non-infiltrated tumors after pembro, with both signatures being mainly driven by the same 8 genes constituted by 3 interferon inducible oligoadenylate synthetase genes (OAS1, 2 and 3, Fig 6E).

MCZ elimination in infiltrated tumors precluded our ability to analyze post-therapy changes in this zone or IZ in this patient group. However, in non-infiltrated tumors T cell proportions increased in multiple neighboring spatial bins near the MCZ core after pembro (Fig 6F). At the subtype level, there were gains in t0 effectors in this same region (Fig 6G). Gene expression changes also supported this subtype change, with CXCL13, GZMB and multiple other genes related to interferon signaling, cytokines, adaptive immune response, and response to pathogens showing upregulation in MCZ T cells from non-infiltrated tumors after pembro (Fig 6H & I). Finally, when we analyzed corresponding scRNAseq + TCR data, we saw almost no TCR expansion changes in infiltrated tumors, whereas most T cell subsets saw expansion gains after therapy in non-infiltrated lesions (Fig 6J).

### Pembrolizumab induces infiltration of CXCL9 expressing myeloid cells in TLS harboring tumors

Several proportional changes outside of T cells were observed in the BCZ of infiltrated and non-infiltrated tumors post-pembro, but these were inconsistent. Infiltrated tumors saw drops in myeloid cell rates in multiple consecutive bins and overall gains in endothelial cells after therapy, while non-infiltrated tumors were reduced for B cells near the periphery (Fig S6A-C). Phenotypically the only pathway changes in BCZ were in non-infiltrated tumors after pembro, when increases in general immune and cytokine related genes were seen (Fig S6D).

In MCZ of non-infiltrated tumors, in addition to the T cell changes reported above, there were reductions of epi. m and endothelial cells near the periphery and overall gains of epi. n and cDC (Fig 7A). This was reflected in shifts away from subniche 1.1 dominated by epi. m. and towards the more heterogeneous subniche 1.0. There were also gene expression changes in non-infiltrated MCZ that were broadly seen across several cell types, including upregulated adaptive immune response, innate immune response, and response to pathogen signals (Fig 7B).

**Figure 7.**
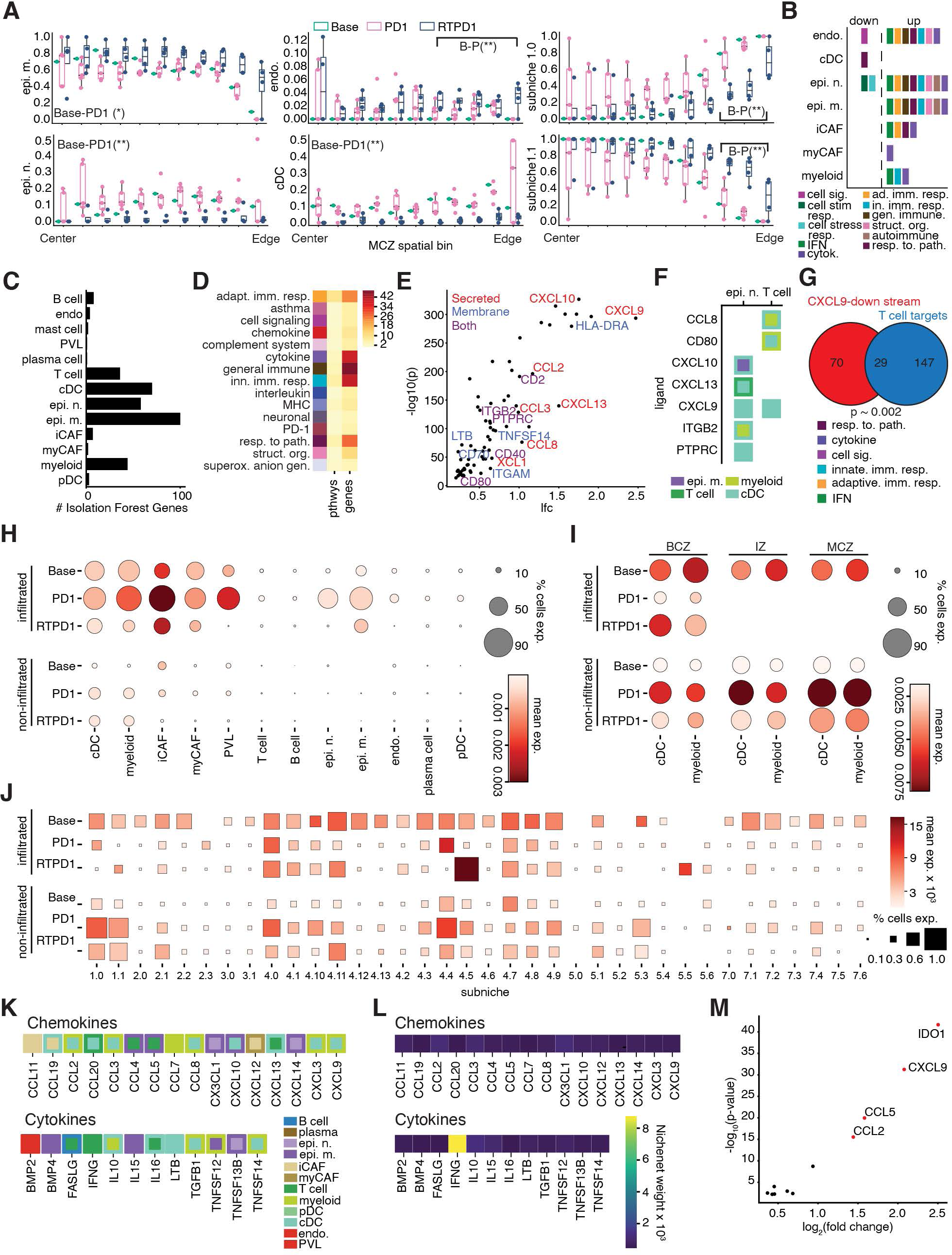
**A.** Sub-niche and cell type proportions through non-infiltrated malignant cell zones which show a statistically significant difference between treatments. Asterisks indicate statistical significance of a Wilcoxon rank sum test between proportions in the bracketed regions between the indicated time points or across the entire zone if no brackets are indicated (* <= 0.1, ** <= 0.05, *** <= 0.01, **** <= 10^-3^, ***** <= 10^-4^). **B.** Summaries of pathways found through differential expression of non-infiltrated malignant cell zones at baseline and PD1. (GSEA; FDR < 0.05). **C.** Bar plot of number of differentially expressed genes in the MCZs (LFC > .2, FDR < .05, and percent expressed in either group > .25) between baseline and PD1 assigned to each cell type by the isolation forest analysis. Note the abundance of pathways in cDCs at PD1. Since cDCs are largely absent in non-infiltrated malignant cell zones at baseline, these pathways were missed in the previous differential expression analysis. **D.** Summaries of pathways describing genes (Enrichr; FDR <= 0.05) brought into the malignant cell zones at PD1 by cDCs (left); Total number of pathways (middle) and genes (right) defining these summaries are indicated in the heatmap. **E.** Scatter plot of the log10 p-value versus log2 fold change of genes brought into malignant cell zones by cDCs after PD1. Secreted ligands are annotated in red. **F.** Ligand-receptor results from cDC secreted ligands in non-infiltrated MCZs. Colors indicate predicted primary (outer) and secondary (inner) senders of ligands (rows) differentially expressed at PD1 compared to baseline anywhere in the MCZs. Ligands shown here explain (NicheNet score > .1) the target genes differentially expressed (LFC > .2, FDR < .05, and % expressed in either group > .25) at PD1 in the target cell types (columns). **G.** Overlap of differentially expressed genes in cDCs at PD1 compared to baseline in the MCZs, and the genes in the highest 10% of weights in the CXCL9 NicheNet table. P-value results from the hypergeometric test for overlap of these two gene sets. Pathway summaries listed are those that describe both gene sets via Enrichr (FDR <= .05). **H.** Dot plots of log-transformed, normalized CXCL9 expression in all cell types in biopsies containing infiltrated and non-infiltrated MCZs in the single cell data. **I.** Dot plots of log-transformed, normalized CXCL9 expression for each zone and infiltration status in cDCs and macrophages. **(J)** Dot plot of cDC and myeloid CXCL9 expression in all subniches. **K.** Upregulated chemokine and cytokine ligands in PD1 MCZs (LFC > .2, FDR < .05, and % expression in either group > .15) where the corresponding receptor is expressed in cDCs (nonzero expression in >30% of cells). Colored boxes show the broad cell type isolation forest results for their expressers. **L.** NicheNet potentials of CXCL9 for each PD1-enriched cytokine and chemokine. **M.** Scatter plot of the -log10 p-value versus log2 fold change of genes comparing cDCs in the non-infiltrated biopsies in the MCZ from PD1 to baseline. The genes included here are those in the top 5% of weights attributed to IFNG in the NicheNet model and considered differentially expressed based on an FDR < .05 and log2 fold change > .2.

To study mechanistically why T cells may have been altered in non-infiltrated MCZ, we calculated overall gene expression changes over time and then assigned the upregulated genes to cells through isolation forests. Though cDC differential gene expression analysis did not indicate pathway changes over time, these cells were amongst the strongest contributors to overall expression changes within MCZ after pembro, likely due to infiltration gains (Fig 7A-7C). Based on the number of supporting genes, beyond general and innate immune response, the most significant gene set brought to the MCZ by cDCs was “cytokine” (Fig 7D). Multiple cDC assigned upregulated genes falling under this annotation were categorized as secreted including CXCL10, CXCL13, CCL2 and CCL8, with the most significantly upregulated being the CXCL9 chemokine (Fig 7E). CXCL9 ligand can recruit T cells and, importantly, ligand-receptor analyses on secreted proteins upregulated in the MCZ found cDC produced CXCL9 best explained the T cell phenotypic change towards effectors in this zone (Fig 7F & G).

CXCL9 is higher in cDC, macs, iCAFS, myCAFs, PVL and epi. m. cells of infiltrated tumors in corresponding baseline scRNAseq data, where therapeutically induced increases in cDC and mac CXCL9 expression in non-infiltrated tumors is also detected (Fig 7H)^25^. The CosMx data also indicated infiltrated tumor macs and cDCs had higher CXCL9 in each TLS associated zone and most subniches at baseline (Fig 7I & J).

However, after pembro CXCL9 myeloid levels increase broadly in non-infiltrated tumors, with the strongest change being in MCZ and corresponding subniches 1.0 and 1.1, where CXCL9^+^ cDCs infiltrate after therapy.

We also searched for ligands that could be causing infiltration of myeloid cells and induction of CXCL9 after pembro in non-infiltrated tumors. PD1 is expressed in both DC subsets and in macrophages, but PD-L1 and PD-L2 are not linked through the nichenet database to CXCL9 (Fig S6F & G). We also studied other cytokines and chemokines that increased in expression in MCZ after therapy (Fig 7K & S6H). Amongst these, IFNG expressed by T cells had the most evidence suggesting it was a modulator of CXCL9, in that CXCL9 falls within the top 5% most highly weighted genes downstream of IFNG in the nichenet database, and in that CXCL9 is the second most upregulated downstream gene of IFNG in myeloid cells (Fig 7L & M).

## Discussion

TLS are well associated with anti-tumor immunity and response to immunotherapy, but questions remain such as: how and why do these ectopic immunological structures form, what drives their variability and what are the implications of that heterogeneity? Prior studies have used tissue staining with limited pools of antibodies, bulk RNA sequencing or spatial transcriptomics to study TLS in pretreatment tumor specimens^3, 24, 32, 33, 34, 35, 36, 37^. Here we used single molecule resolution spatial transcriptomics to study longitudinal biopsies taken throughout an immunotherapy response. This allowed cell type specific and spatially defined molecular programs of response to be elucidated and for a comprehensive list of molecular level spatial interactions involved in an immunotherapy response to be drafted.

Our computational framework for defining and examining TLS and associated structures allowed identification of two tumor groups based on malignant cell zone cellular proportions. One group displayed higher baseline signals of immunoreactivity in all cell types within the MCZ and had higher myeloid and T cell MCZ infiltration.

Coincidently, this group had higher baseline T cell expansion rates and tumor bed elimination after pembro alone. The other group only displayed MCZ T cell infiltration after pembro and tumor bed elimination after pembro with radiation and was broadly characterized by inflammatory fibroblasts expressing collagen signals. Prior work on TLS heterogeneity showed mature TLS with segregated B and T cell zones are associated with therapeutic response and long-term survival. Notably, “infiltrated” tumors have better separation of B and T cell zones and their intermediate zones between TLS and tumor beds also show higher immunoreactivity, providing evidence TLS function directly regulates tumor bed infiltration and elimination.

One explanation for the observed baseline infiltration differences was found when searching for gene programs whose spatial patterns within structures differed between tumor groups. We found MHC expression was similar in infiltrated and non-infiltrated tumors at the periphery of tumor zones, but that malignant cell MHC expression rose dramatically only in the infiltrated lesions moving towards the tumor core. This indicates TLS maturity and function can be guided by antigen availability within nearby tumor beds.

We were unable to compare tumor beds or intermediate zone response dynamics between infiltrated and non-infiltrated tumors, due to clearance in infiltrated tumors after therapy. However, in non-infiltrated tumors T cells increased in MCZ after therapy where they also transitioned towards effectors, coincident with overall TCR expansion gains. At the same time T cells became more prominent at TLS, indicative of the structures playing a role in T cell entry to the site after therapy. Further evidence for this comes from naive cells being more prominent at infiltrated TLS prior to therapy and the fact that naive signatures are prominent in the T cells found at non-infiltrated tumor TLS after pembro.

Though the cDCs that gain entry to non-infiltrated MCZ after therapy are not detected to increase at the TLS concurrently, these cells were enriched in intermediate zones at baseline. This suggests the local tissue environment of the TLS, even if it does not extend into the tumor bed, plays a role in tumor clearance. Though multiple ligand-receptor analyses were performed, the most relevant finding was increased CXCL9 brought by cDCs infiltrating the non-infiltrated tumoral regions after pembro. CXCL9 is a predicted ligand driver of the effector T cell phenotype that arises coincidently. Notably, CXCL9 was higher in expression tumor wide in infiltrated tumors prior to therapy, suggesting this molecule is important for differentiating tumor elimination potentials of TLS harboring tumors.

## Supplementary Figure Legends

**Figure S1.**
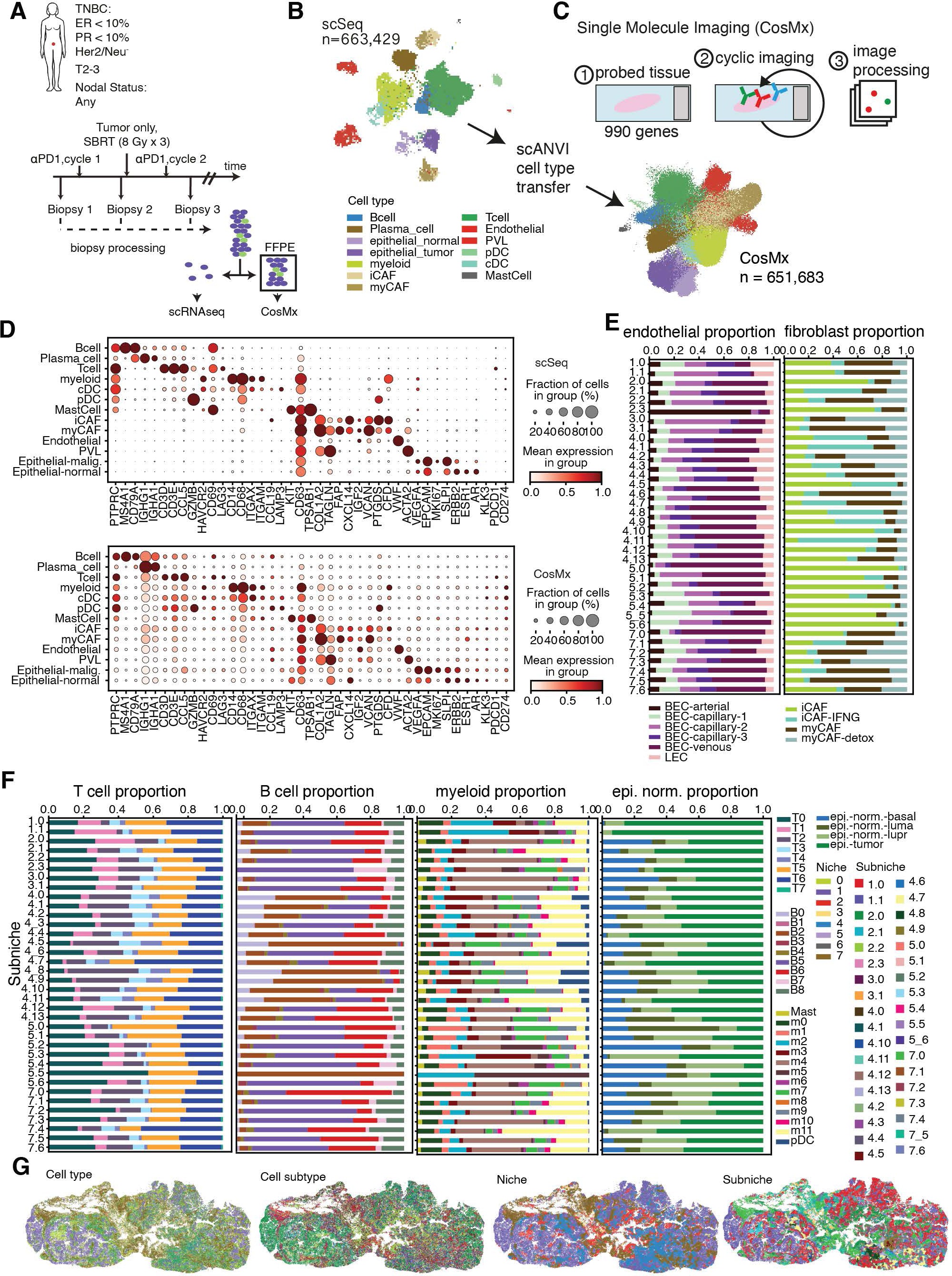
**A.** Schematic of clinical trial design for single cell and CosMx data collection. **B.** UMAP of cell type annotations in the scRNA-seq data. **C.** Schematic for CosMx experimental design (top) and UMAP of cell type classifications in the CosMx data that were obtained from scANVI label transfer from scRNA-seq cell type annotations (bottom). **D.** Dot plot of canonical marker gene expression for each cell type in the scRNA-seq data (top) and CosMx data (bottom). Size of each dot indicates percentage of cells expressing each gene, and the color intensity correlates with the standard-scaled, average log-normalized expression. **E.** Stacked bar plots of the proportion of all cells in each subniche belonging to each cell subtype for endothelial cells and fibroblasts along with **F.** T cells, B cells, myeloid cells, and epithelial cells. **G.** Spatial locations of cells in biopsy h02A colored by cell type, cell subtype, nice, and subniche, respectively.

**Figure S2.**
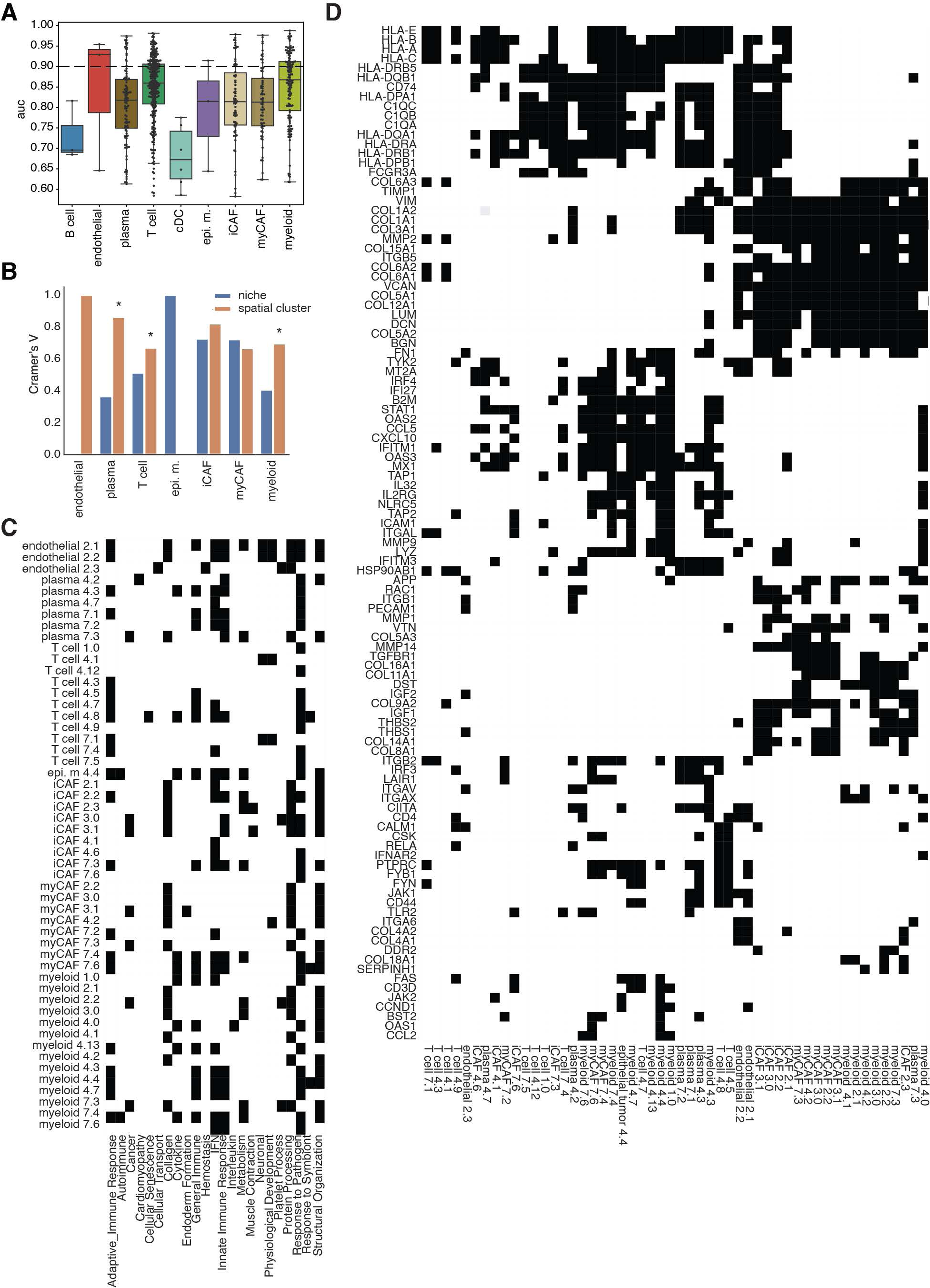
**A.** Box plots of AUC for predicting subniche membership between two subniches for each cell type. Dashed line indicates AUC cutoff for subniche comparisons in which cell types are deemed to have transcriptionally distinct profiles. **B.** Bar plot of Cramer’s V effect size comparing for every cell type, clustering of Jaccard scores for pairwise subniche gene sets to spatial clusters (orange) and broad niche groupings (blue). Asterisks indicate a chi-squared contingency test p-value < .05. **C.** Pathway summaries for the target gene sets used in the ligand-receptor analysis in Fig. 2. **D.** Pathway genes defining the previous summaries. Here we show only the 30% most common genes in this set of pathways.

**Figure S3.**
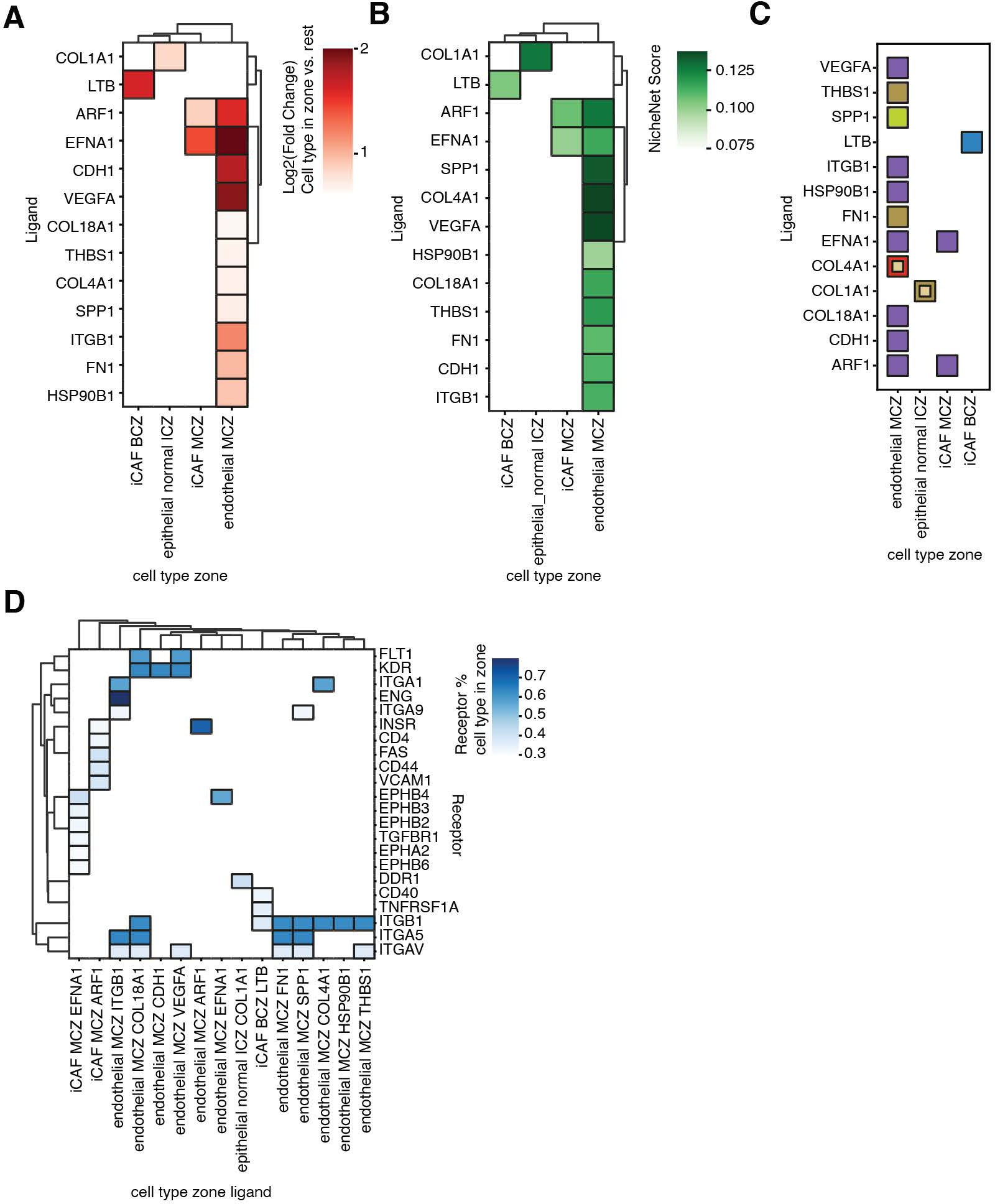
**A.** Heatmap of Log2-fold change of interacting ligands in each cell type between each zone and the rest of the zones for BCZ, ICZ, and MCZ, and between TCZ and BCZ. **B.** Heatmap of NicheNet scores between each interacting ligand and each target condition’s (zone and cell type combination) target gene set. **C.** Predicted primary (outer) and secondary (inner) senders of each interacting ligand and each target condition. **D.** Percentage of cells in each target condition expressing receptors associated with interacting ligands.

**Figure S4.**
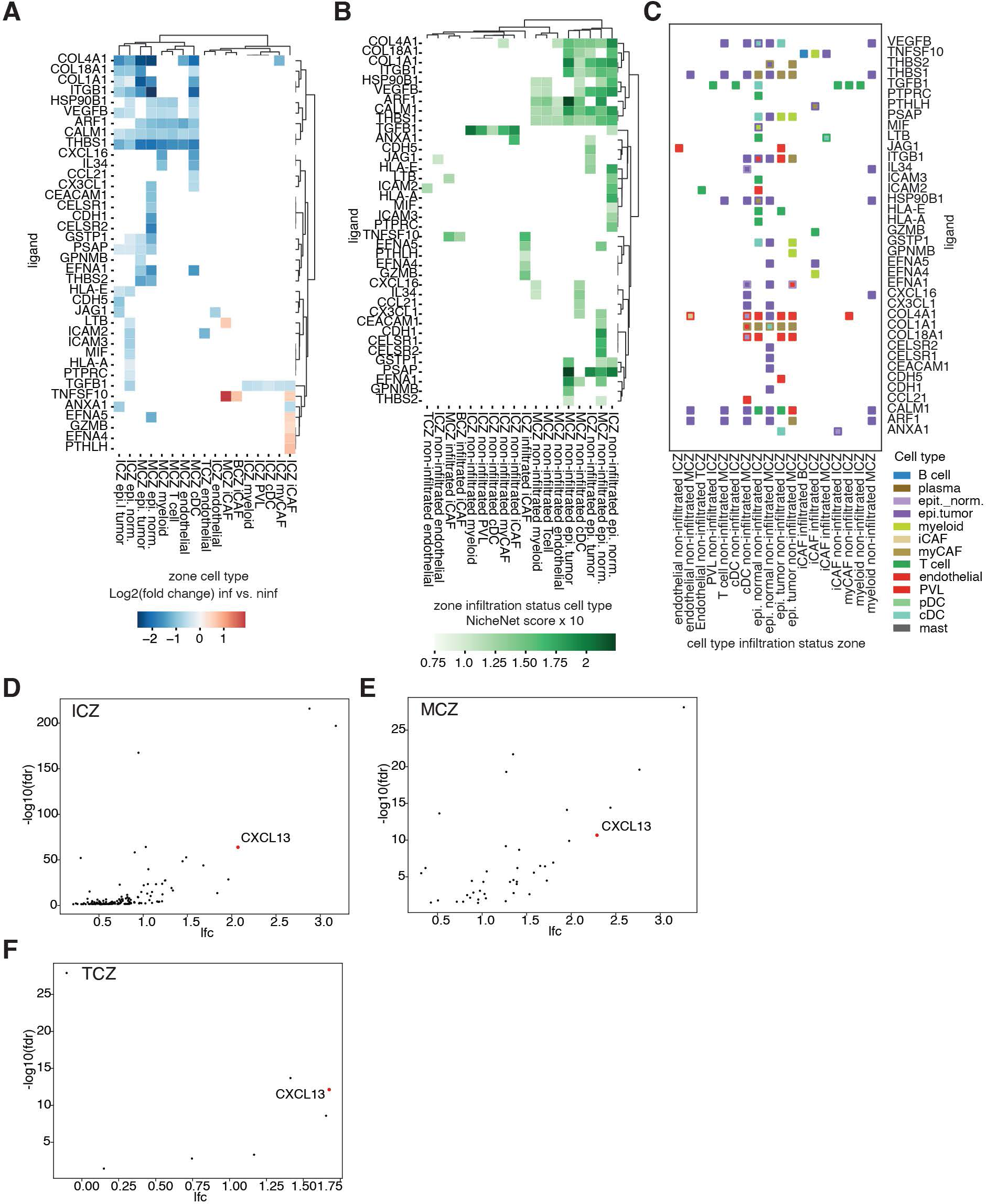
**A.** Heatmap of Log2-fold change of interacting ligands in each cell type and each zone between infiltrated and non-infiltrated biopsies. **B.** Heatmap of NicheNet scores between each interacting ligand and each target condition’s (zone, infiltration status, and cell type combination) target gene set. **C.** Predicted primary (outer) and secondary (inner) senders of each interacting ligand in each target condition. **D.** Volcano plot of genes upregulated in the T cells of non-infiltrated tumors after pembrolizumab in BCZ. **E.** Volcano plot of genes upregulated in the T cells of non-infiltrated tumors after pembrolizumab in MCZ. **F.** Volcano plot of genes upregulated in the T cells of non-infiltrated tumors after pembrolizumab in TCZ.

**Figure S5.**
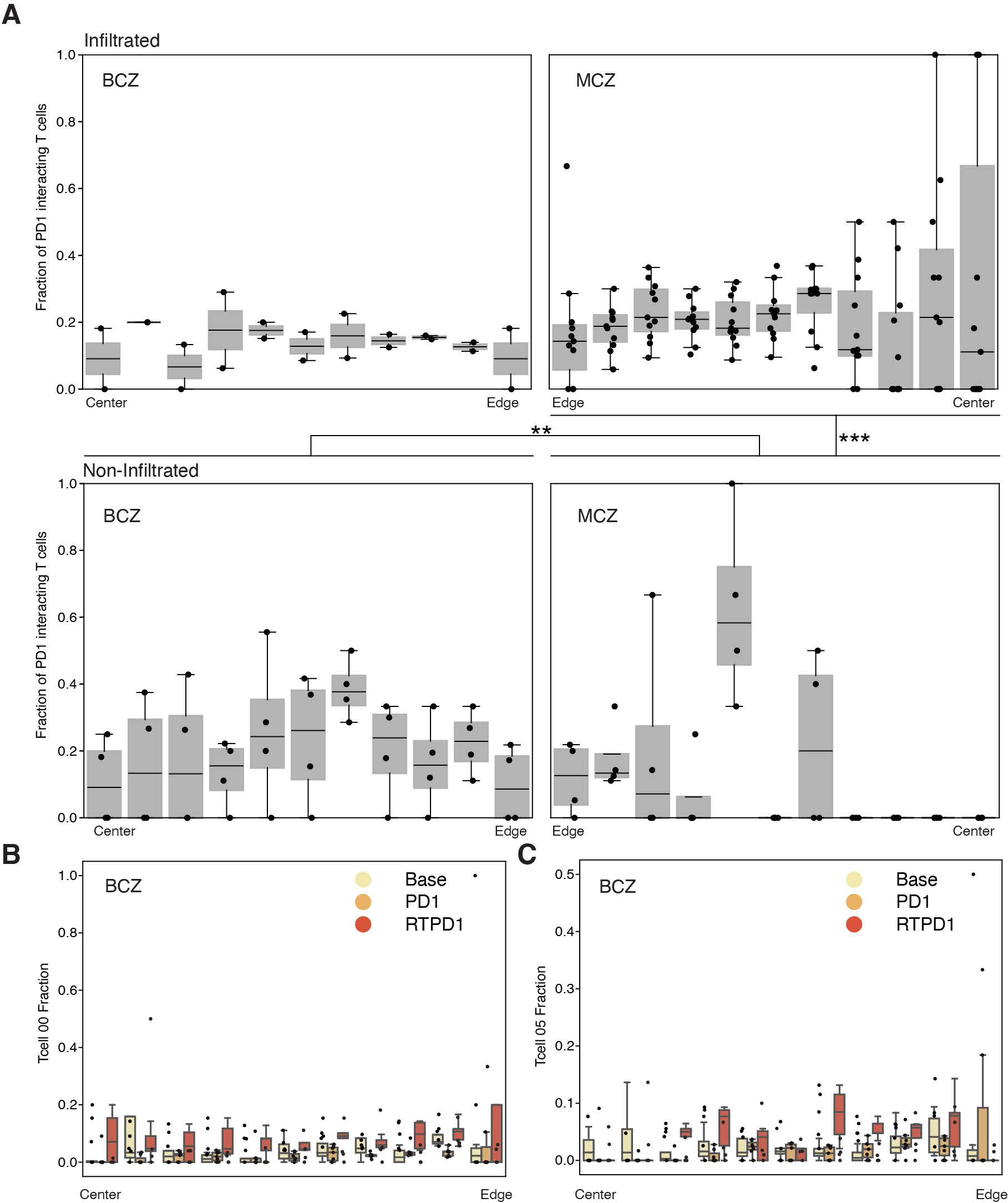
**A.** Box plots of proportions of PD1 interacting T cells in the statistically different BCZ and MCZ at baseline in infiltrated tumors (top) and non-infiltrated tumors (bottom). Asterisks indicate statistical significance of a Wilcoxon rank sum test between proportions in the bracketed regions (* <= 0.1, ** <= 0.05, *** <= 0.01, **** <= 10^-3^, ***** <= 10^-4^). **B.** Box plots of T cell subtype 0 proportions moving through the BCZ at all time points in non-infiltrated tumors. Asterisks indicate statistical significance of a Wilcoxon rank sum test between proportions across the entire zone in the indicated time points (* <= 0.1, ** <= 0.05, *** <= 0.01, **** <= 10^-3^, ***** <= 10^-4^). **C.** Box plots of T cell subtype 5 proportions moving through the BCZ at all time points in non-infiltrated tumors. Asterisks indicate statistical significance of a Wilcoxon rank sum test between proportions across the entire zone in the indicated time points (* <= 0.1, ** <= 0.05, *** <= 0.01, **** <= 10^-3^, ***** <= 10^-4^).

**Figure S6.**
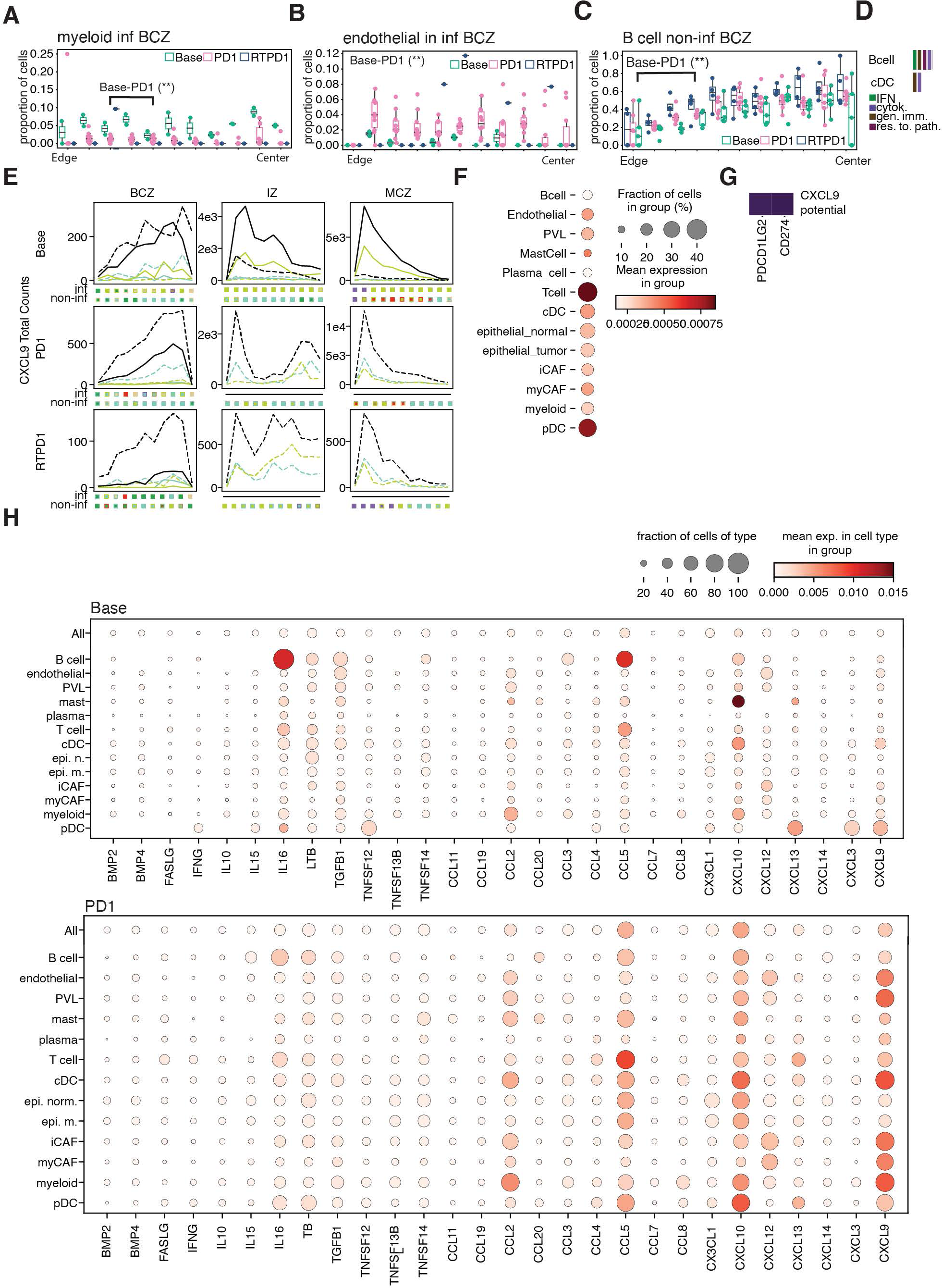
**A.** Myeloid cell proportions in infiltrated BCZs at all treatments. Asterisks indicate statistical significance of a Wilcoxon rank sum test between proportions in the bracketed regions (* <= 0.1, ** <= 0.05, *** <= 0.01, **** <= 10^-3^, ***** <= 10^-4^) **B.** Endothelial cell proportions in infiltrated BCZs at all treatments. Asterisks indicate statistical significance of a Wilcoxon rank sum test between proportions in the entire region where a bracket is not indicated (* <= 0.1, ** <= 0.05, *** <= 0.01, **** <= 10^-3^, ***** <= 10^-4^) **C.** B cell proportions in non-infiltrated BCZs at all treatments along with upregulated pathways in these regions. Asterisks indicate statistical significance of a Wilcoxon rank sum test between proportions in the bracketed regions (* <= 0.1, ** <= 0.05, *** <= 0.01, **** <= 10^-3^, ***** <= 10^-4^) **D.** Pathway summaries for pathways describing genes upregulated in B cells and cDC in BCZ of non-infiltrated tumors. **E.** CXCL9 counts moving through each zone at each time point in all cells, cDCs, and myeloids. Results of isolation forest identification of the CXCL9 expressing cell type is shown in the bottom panel. **F.** Dot plot of PD1 gene expression across all cell types after pembrolizumab treatment in non-infiltrated MCZs. **G.** Heatmap of CXCL9 NicheNet downstream weights for PDL1 and PDL2. **H.** Dot plot of expression of cytokines and chemokines identified in Fig. 7K in non-infiltrated MCZs at baseline and after pembrolizumab.

## Methods

### scRNA-seq data preprocessing/cell typing

Aligned and mapped scRNA-seq data were individually subjected to quality control filtering using Scanpy (Wolf, et al., 2018) and Scrublet (Wolock et al., 2019). Initial filtering was performed to remove cells with >10% of their counts mapping to mitochondrial genes, and to remove cells with fewer than 100 unique mapped genes. Further, the top 1% of genes by counts from each library were removed as potential doublet UMIs. Next, the doublet prediction tool Scrublet was applied using the top 10% of highly variable genes. A threshold of 0.25 on the predicted doublet score was used to remove potential heterotypic doublet cells.

Following initial quality control, cells from all libraries were concatenated and clustered at once using single cell Variational Inference (scVI; Lopez et al., 2018). The scVI latent space was used for clustering with the leiden algorithm and was used to generate a UMAP representation of the learned latent space (RAPIDS.ai cuML, cuGraph). Clusters were annotated as each of the major immune and tumor microenvironment component cell types including fibroblasts, cancer associated fibroblasts (CAF) and epithelial cells based on expression of canonical marker genes.

Copy number variant profiles were estimated for each epithelial cell using inferCNV [https://github.com/broadinstitute/inferCNV]. All cells from each non-epithelial cell type (’Tcell’, ’myeloid’, ’myCAF’, ’Plasma_cell’, ’Endothelial’, ’Bcell’, ’cDC’, ’iCAF’, ’PVL’, ’pDC’, ’MastCell’) were used as reference. Epithelial cells from individual patients were encoded as "Epithelial_{patient}" to allow grouping of malignant cells by patient. "infercnv::run" from infercnv (v 1.13.0) was ran with the following parameters: "cutoff": 0.1, "cluster_by_groups": FALSE, "denoise": TRUE, "HMM": TRUE, "cluster_references": TRUE, "analysis_mode": ’samples’, "z_score_filter": 0.

CNV states given by HMM i6 model and filtered based on posterior probability using a Bayesian network were used to identify cancer cells. Briefly, for each epithelial cell, we calculated the percentage of genes showing copy loss or gain (CNV state not equal to 3) and assigned a percentage mutation metric. For each epithelial sub cluster identified previously, we calculated the median percentage mutation across all cells and assigned each cluster a normal/cancer annotation based on the z-score.

Endothelial, CAF and epithelial cells were further subtyped by performing scVI a second time on cells of each major cell type. Immune cells were assigned subtypes based on previous annotations of the immune cells from CD45 enriched libraries in a previous study. To systematically transfer subtype annotations, single-cell ANnotation using Variational Inference (scANVI; Xu et al., 2021) was applied to cells of each broad immune cell type (Bcell, Tcell, Myeloid) individually.

### CosMx data preprocessing

All quality control was run on a per biopsy basis. Cells were filtered out if they had fewer than 20 gene counts, expressed fewer than five genes, or expressed greater than the 99.9^th^ percentile of counts across all cells. Additionally, we utilized the negative probe gene expression data to home in on signal from biological probe expression. For every cell, we computed the median count across all negative probes. Any gene in a cell whose expression surpassed the median negative probe expression was set to 0. For most downstream analyses, gene expression was normalized by the total expression in each cell and scaled by a factor of 500 using the Scanpy function normalize_total(), and natural log-transformed using the Scanpy function log1p() (v. 1.9.3).

### Classifying cell types and subtypes in the CosMx data

We utilized single-cell ANnotation using Variational Inference (scANVI; scvi-tools v. 0.14.5; Xu et al., 2021) in order to transfer previously defined cell labels from the matched scRNA-seq data set (described above) to our CosMx data. A set of 822 genes were curated by filtering lowly and highly expressed genes and intersecting the remaining genes with the scRNA gene set. Latent dimensions were first learned on the scRNA-seq raw gene counts using single cell Variational Inference (scVI; scvi-tools v. 0.14.5; Yosef et al., 2022). The model was formulated with 64 latent variables and 5 hidden layers and allowed to train over at most 400 epochs (early_stopping = True). The scANVI model was then initialized with the scRNA-seq reference data and pre-trained scVI model. Training occurred over 100 epochs, and with n_samples_per_label set to 100. The scANVI model was then trained on the CosMx raw gene counts over a maximum of 100 epochs (early_stopping = True; weight_decay = 0). Lastly, the model predicted cell types on the CosMx query cells (Fig S1C & D).

For cell subtype classification, we split our CosMx data based on five broad cell types: fibroblasts (iCAFs and myCAFs), endothelial cells, normal epithelial cells, lymphoid cells (B cells, plasma cells, and T cells), and myeloid cells (cDCs and macrophages). Subtype annotations from each broad cell type category in the scRNA-seq data set were transferred to the CosMx data using the same protocol outlined for broad cell type labeling. For fibroblasts, endothelial cells, normal epithelial cells, and lymphoid cells, broad cell types were modified by inheriting the broad cell type class of the new cell subtype predictions.

UMAP representations for all cells and each broad cell type subcategory were generated by running first, Scanpy’s neighbors() (v. 1.9.3) function on the scANVI latent dimensions and second, Scanpy’s umap() function on these batch-corrected neighborhoods.

### Tier 1 Niching

In order to describe recurring cell type neighborhoods across our multiple biopsies, we implemented a custom analysis for “niche” detection. We first computed the average radius of a cell by taking the average segmentation area across all cells, A, and calculating the ceiling of the square root of A/pi. For every cell, we defined its local neighborhood to be any cell within an approximate radius of two cells using scikit-learn’s radius_neighbors_graph() method (scikit-learn v. 1.2.1; Pedregosa et al., 2011). We then defined a matrix describing the percentage of each cell’s local neighborhood that belonged to each broad cell type class. An adjacency matrix was computed using cuML’s kneighbors_graph() method, drawing connections between similar cell neighborhood profiles, and was provided as input for cuGraph’s leiden() implementation (Rapids.ai v. 22.02). We ran this framework using all combinations of the following parameters: Leiden resolution = {0.2, 0.4, 0.6, 0.8, 1.0, 1.2, 1.4, 1.6}, k for kneighbors_graph() = {20, 30, 60}, and distance metric for kneighbors_graph() = {euclidean, cosine}. For all runs of Leiden clustering, we computed the silhouette score using scikit-learn’s silhouette_score() (v. 1.2.1) function, specifying sample_size = 10000 and random_state = 0. We ultimately used the niches defined from Leiden clustering utilizing the parameters that resulted in the highest silhouette score.

Upon evaluating the proportion of cells in each niche belonging to each cell type class, we noticed sets of niches that were highly correlated to each other. In order to find a more distinct set of niches based on cell type composition, we conducted an iterative approach for merging highly correlated niches. In each iteration, so long as there were pairs of niches whose cell type compositions correlated beyond a correlation coefficient of .95, we merged the pair of niches with the highest correlation coefficient by computing their average cell type composition.

For cells that lacked a cellular neighborhood within a radius of 2 cells, we post factum assigned them the most prominent niche amongst their 50 nearest neighbors (scikit-learn v. 1.2.1); NearestNeighbors()).

### Tier 2 Niching

In order to obtain more granular niches, we defined “subniches” by considering recurring cell subtype neighborhoods (Fig 1 & S1E, F). Here, local neighborhoods were defined by taking the Delaunay triangulation of cell locations in each biopsy, utilizing scipy’s Delaunay method (v. 1.7.1), and removing any edges with a distance > 500 pixels. For every cell, we performed a breadth-first search on the Delaunay graph using NetworkX’s bfs_successors() (v. 2.5.1) method, and found all cells within a maximum search depth of 4. The neighborhood matrix was defined by computing the proportion of cells in each neighborhood belonging to each subtype class. For every broad niche, excluding niches 0 and 6 due to low cell numbers, and for all values of k between 2 and 30, we used scikit-learn’s (v. 1.0.1) MiniBatchKMeans() method (batch_size = 2048; random_state = 999) for clustering the subtype neighborhood profiles. For every run of clustering, the average silhouette score (scikit-learn silhouette_score(); v. 1.0.1) was computed across 20 samples of 10,000 data points. The subniche clustering resulting from the value of k that produced the highest average silhouette score was used. Subniches with fewer than 2500 cells were flagged and set aside in further analyses.

### Defining ligand interactions in each subniche at baseline

First, we define a graph drawing connections between nodes representing cells of different cell types and subniches. The starting set of nodes was all cell types in all subniche contexts. Nodes were then filtered if they represented less than 2500 cells across all four patients, h12, h43, h02, and h16, and if their cell type comprised less than 10% of its subniche’s cells. Shift edges were defined to be mappings between groups of cells of the same type in two different subniches that have the potential to show spatially-influenced phenotypic differences. For a given cell type in a subniche, pressure edges connect cells of a different type that have the potential to affect the phenotypic shift of the cell type from the starting subniche to another.

We ran a protocol adapted from Augur (Skinnider et al., 2020) in order to determine if there were transcriptional differences between shift edges. We first defined the feature space to be an scVI-derived latent space consisting of 32 features, generated by specifying 3 layers with 128 units, running over a maximum of 300 epochs and flagged for early stopping. For every subniche pair combination in every cell type, if available, we took 100 subsamples of 200 cells from both populations using numpy’s random.choice() method (v. 1.20.3). For every subsample, a ridge classifier model was trained on a set of cells the size of 30% of the population (scikit-learn’s train_test_split(); v. 1.0.1). The trained model was then used to predict each subniche class amongst the test population, and the AUC score (scikit-learn’s roc_auc_score(); v. 1.0.1) was recorded using the model’s decision function scores.

We then employed “pruning” of our graph based on cell numbers, Augur scores, and spatial correlations. Any node with fewer than 1000 cells were removed. Any cell type in a subniche that was classified with an average AUC > .9 in comparison to the same cell type in another subniche was maintained. If any pressure edge did not have any downstream shift edges, it was removed. Additionally, we require pressure edges to occur between nodes that are spatially correlated with one another.

All remaining pressure-to-shift node triplets were further interrogated with our ligand-receptor analysis. Differential expression was conducted on all passing cell type/subniche pair comparisons using scanpy’s (v. 1.8.2) Wilcoxon test. For every comparison, we defined the set of target genes for each receiving population to be those that passed the following thresholds: absolute value of the log2 fold change > .5 and FDR < .1. We moved forward with the receiving population’s target gene set if it contained at least 20 genes.

We next defined plausible sets of receptors in the receiving cell populations and corresponding ligands that might bind to the receptors and affect downstream target gene expression. To do so, we found all receptors that are expressed in at least 20% of each target cell population. Available ligands were defined as those expressed in at least 20% of any pressure node leading to the target cell population. Additionally, the ligands were required to be maximally expressed in the sending node compared to all other nodes containing the sending node’s cell type. We then adapted the scoring mechanism from NicheNet in order to determine to what extent each plausible ligand is known to affect the downstream perturbed genes in the target population. That is, for every plausible ligand affecting the target cell population’s gene expression program, we pulled the NicheNet-defined vector of weights associating the ligand to each of the genes belonging to our CosMx gene panel. We defined a binary vector, where for every gene in the background set, 1 indicated if the gene belonged to the target gene set, and 0 otherwise. Pearsonr correlation was run to associate the NicheNet weight vector with the binary vector, giving us a score that indicates the strength of association between each ligand and the target gene set. We defined our final filtered set of ligands associated with the target cells’ gene program to be those achieving a NicheNet score > .1.

Lastly, we looked for cell type populations acting as the most likely senders of each ligand found to be associated with each target population. Only cell types that expressed the ligand in 2.5% of cells in the matched single cell data set, achieved mean nonzero expression of at least .1, and whose population exceeded 30 cells were considered. If only one cell type passed these thresholds, this cell type was called the sender. Otherwise, we ran the Isolation Forest anomaly detection method (scikit-learn v. 1.2.1) on two metrics for ligand expression for each of the putative sending cell types: mean nonzero expression and total expression. We computed the z-score of each feature across all the cell types. Any cell type that was not only called an anomaly, but also achieved scaled features > 0 was called a sending population. If no cell type met these criteria, we computed a pseudo anomaly threshold. Namely, we identified cell types that not only had two positive scaled features, but also had an anomaly score < .5 times the average anomaly score. If no true anomalies or pseudo anomalies were detected, we identified the sending populations to be those with the top two highest average nonzero expression values.

### Correlation of cell type/subniche phenotypes to spatial clusters and broad niches at baseline

For each cell type, we computed the Jaccard score between sets of genes defining every subniche pair. We then used scikit-learn’s (v. 1.2.1) AgglomerativeClustering() method, with Ward’s method for computing linkages, in order to define clusters of subniches in which a cell type has similar gene expression programs. In order to determine the number of clusters, we iteratively ran agglomerative clustering specifying “n_clusters” to be 4 through the maximum number of gene signatures for any cell type, and computed the silhouette score for each set of clusters generated. The value of “n_clusters” that achieved the maximum silhouette score was ultimately used for cluster generation. We then compared clusters of transcriptionally similar subniches for each cell type to each subniche’s broad niche category, and to each subniche’s spatial clustering. To do so, we ran the Chi2 contingency test to determine whether or not there was statistically significant overlap between each comparison of labels, and computed Cramer’s V to measure the effect size of this overlap.

### Architectural analysis

In order to quantify the spatial clustering of cells of different types, niches, and sub-niches we compute the mean spatial cross correlation coefficient, w, for each categorical pair a,b by solving the equation:

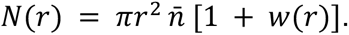

Here, N(r) is the number of cells of type b within a circle of radius r around a cell of type a. *n̄* is the average density of cells of type b across the entire sample. Therefore, the cross correlation coefficient categories the over- or under-density of cells of type b, relative to the background density, within a scale r of cells of type a.

We compute this for all cell type, niche, and sub-niche pairs, and we will use the sub-niche clustering results throughout this work to define tissue architecture. Spatially associated clusters are determined by clustering the vector of sub-niche correlations at a fixed scale of 500 coordinate units. This scale was chosen as the typical scale at which positive correlations peak. To determine the statistical significance of each spatial cluster we performed a bootstrap analysis, recomputing clusters in each of 10,000 samples bootstrapped at the scale of individual FOVs. For each sub-niche, we then computed the average rate at which it clustered with other sub-niches in its baseline cluster.

### Determination of Regions of Interest

Throughout this work we refer to four regions of interest: the B cell zone at the core of a TLS, the T cell zone extending from this core, the malignant cell zone, and the intermediate zone between the B cell and malignant cell zones. The B cell zone was simply defined by the sub-niches with the highest proportion of B cells. The B cell density field is highly concentrated in local peaks which typically correspond to connected spatial clusters of sub-niches 4_5 and 4_9. These two sub-niches were found to have a very high cross correlation coefficient on small scales. Sub-niche 4_8 is also highly spatially correlated with these two sub-niches and contains a very high proportion of T cells. This sub-niche tends to extend away from the B cell core, and we define it as the T cell zone of the TLS. The malignant cell zone is defined by Niche 1 which contains the highest proportion of epithelial tumor cells. Typically these clusters of cells are structured with a dense core of tumor cells in sub-niche 1_1 surrounded by a border of sub-niche 1_0.

For much of our further analysis we will define a directionality to each zone in order to understand changes in the tissue as you move through a region. To do this we start with an image of an individual instance of one region. We then convolve this image with a gaussian filter of a size chosen to smooth over any small holes in the structure of interest. We then compute the gradient of this image and interpret the resulting vector field as pointing towards the center (or, for more complex structures, something akin to the center of mass) of the zone.

With these three zones and their directionality we then define an intermediate zone stretching from the B cell zone to nearby malignant cell zones. We compute the angle between the vector field pointing toward a B cell zone with those pointing towards nearby malignant cell zones and define the intermediate zone as any region where the angle between the two vectors is greater than 0.8π radians.

### GSEA Analysis

We use the Gene Set Enrichment Analysis (GSEA) tool (gseapy v. 1.0.4) to determine biological pathways which describe sets of differentially expressed genes throughout this work. Using Scanpy (v. 1.9.2), we first compute a ranked list of genes based on differential expression between two subsets of a cell type population (Scanpy wilcoxon v. 1.9.2). We apply cuts for significant differences, ensuring that each gene is expressed in at least 2.5% of that cell type in the matched scRNA data, expressed in at least 25% of cells in one of the populations, has an FDR of less than 5%, and a log fold change of at least 0.2 in either direction. Finally, the log2 fold changes between the two cell populations of the filtered gene set was passed to GSEA and compared against the KEGG 2021, GO Biological Processes 2021, and Reactome 2022 human databases.

We performed a similar analysis for gene expression programs defined by our gradient analysis. In this case, the ranked list of genes provided to GSEA was simply the list of mean dot products with the structural vectors defining each region.

To report these pathways we grouped them into a collection of summarized categories in order to avoid overlapping pathways from multiple databases. We also clustered the leading edge genes from GSEA to find groups of pathways defined by the same sets of genes using the sklearn (v. 1.2.0) AgglomerativeClustering method (S2D).

### Gradient Analysis

We implemented a new gradient analysis to compute spatial changes in gene expression across structures in the tissue. First, we convolved images of the expression of each gene in each celltype with a gaussian filter with a radius of 200 coordinate units. We then took the gradient of this field and divided by the local density of that cell type in order to get the expression gradient per cell. Finally, we took the dot product of this vector field with the structural vectors defined above for each structure to determine the component of expression moving toward or away from the core of each zone, or from the B cell zone to the malignant cell zone in the case of the intermediate zones. We compared the resulting distribution of gradients across each zone to a uniform angle distribution, performing a KS test to ensure a statistically significant detection of a gradient in a direction of interest.

### Defining ligand interactions in each spatial zone at baseline

First, we defined our target cells, or cells that potentially receive a ligand signal, and sending cells, or cells that may send a ligand signal to the target cells. In the case where we looked for ligand interactions defining cell populations in each spatial zone (i.e., malignant cell zone, B cell zone, T cell zone and intermediate cell zone) at baseline, the target cell populations were cells of each cell type in each zone at baseline. The sending populations of each target cell set were found using the Delaunay triangulation of cell locations in each biopsy, where for every target cell, we performed a breadth first search using NetworkX’s (v. 2.5.1) bfs_successors() method, and found all cells within a maximum search depth of 2.

Next, we defined target genes that were enriched in the target cell populations, and ligands that were enriched in the sending cell populations. Target gene sets in the target cells in the malignant cell zone, intermediate, and B cell zones were defined by Scanpy’s (v. 1.9.3) rank_genes_groups() function, specifying the Wilcoxon test, and comparing for every cell type, gene expression between cells in each zone and cells in the rest of the zones. For the T cell zone, differential expression was run on every cell type, comparing cells in the T cell zone with cells in the B cell zone only. Differential expression was run only if the number of cells in all zones being compared exceeded 30. Target genes that achieved an FDR < .05, were expressed in at least 30% of either of the two populations, surpassed a log2 fold change > .5, and were expressed in at least 2.5% of the target cell type population in the matched scRNA-seq data set were maintained. Furthermore, target gene sets were required to be at least ten genes. Ligands potentially affecting the target cells’ gene expression programs were determined by running the same comparisons on the neighboring cell populations and pulling differentially expressed genes overlapping with NicheNet’s curated list of ligands. Ligands were filtered based on the same FDR, log2 fold change and expression rate thresholds.

We further filtered ligands using a series of ligand prioritization steps. First, we utilized NicheNet’s algorithm for associating ligands with target gene sets. The following algorithm was run for each cell type in each spatial zone: For a given cell type in a spatial zone, we defined the background set of genes to be the union of genes found in the target gene set and all genes expressed in at least 10% of the cells. For every ligand found to be enriched in the neighboring sending population, we used NicheNet’s scoring algorithm (defined above) to associate ligands to the target gene set. Ligands achieving a score > .1 were maintained. Additionally, we filtered for ligands associated with at least one receptor (based on NicheNet’s database) that was expressed in at least 30% of the target cells. The same anomaly detection protocol defined previously for sender cell type assignment was used.

### Defining ligand interactions across infiltration status and treatment

In order to define interactions that differentiate cells in each cell type and zone between infiltration status at baseline, we queried target cells of each cell type, spatial zone, and infiltration status at baseline. Here, target gene signatures were defined by the Wilcoxon test (Scanpy’s rank_genes_groups() method) comparing for every cell type in each spatial zone, cells in infiltrated biopsies with cells in non-infiltrated biopsies at baseline. Putative ligands associated with target gene expression were found by running the same comparisons for the neighborhoods of each target cell population. All FDR, log2 fold change, and expression rate cutoffs for differential gene expression, as well as all ligand prioritization thresholding were the same as those used when comparing cell types across spatial zones at baseline. In order to add confidence in ligand interactions that occurred broadly across multiple structural instances, we implemented the additional cutoffs: for every prioritized ligand and associated target gene set enriched in a cell type/spatial zone in biopsies of a certain infiltration status, we ensured that the median average ligand expression in the neighboring cells of targets across structural instances was greater in the infiltration status of interest compared to the opposing infiltration status. Additionally, we checked that the median average target gene signature in the targets across structural instances was greater in the infiltration status of interest compared to the opposing infiltration status. In order to determine the most-likely senders of each ligand, we used the same anomaly detection methodology as before.

Lastly, we looked for interactions that were enriched after each treatment in each cell type, zone and infiltration status. The target gene sets for each target cell population were defined as follows: Differential expression (Scanpy’s Wilcoxon test with rank_genes_group()) was conducted for every cell type and infiltration status between Base and PD1, and PD1 and RTPD1. Tests were only run if the number of cells in both groups exceeded 15. In B and T cell zones, for every cell type and treatment (in each comparison), genes enriched in one infiltration status and not the other were used as target genes. For malignant cell and intermediate cell zones, target genes were defined as those enriched in every cell type and treatment per comparison in non-infiltrated biopsies. The same comparisons were made for the cell neighborhoods surrounding every target cell set and identifying differentially expressed ligands from NicheNet’s list. Genes were considered enriched in each comparison if they achieved an FDR < .05, were expressed in at least 25% of either of the two populations, surpassed a log2 fold change > .2, and were expressed in at least 2.5% of the target cell type population in the matched scRNA-seq data set. A target gene set was only considered moving forward if it contained at least ten genes. All ligand prioritization thresholds were the same as those used in previous ligand interaction analyses. Similar to what was done in identifying ligand interactions defining infiltration status in each cell type at baseline, we looked for ligand interactions specific to infiltration and treatment status that were enriched across multiple instances of each spatial zone. To do so, for every prioritized ligand and associated target gene set enriched in a cell type in biopsies of a certain infiltration status and treatment, we ensured that the median average ligand expression in the neighboring cells of targets across structural instances was greater in the treatment of interest compared to the opposing treatment. Additionally, we checked that the median average target gene signature in the targets across structural instances was greater in the treatment of interest compared to the opposing treatment.

### Ligand interaction analysis on gene gradients

In order to find ligands interacting with target genes based on shared directionality across each spatial zone at baseline, we conducted the following protocol: For each cell type in each zone at baseline, we defined positive gradient signatures as the genes passing the 95th percentile of positive dot products with respect to the principal axis of a zone, and negative gene gradients as the genes less than the 5th percentile of negative dot products. Additionally, for every gene set, genes were required to be expressed in at least 30% of the cell population in the CosMx data, and 2.5% of the cells’ cell type population in the scRNA-seq data. Moving forward, we only considered target gene sets containing at least ten genes. For each pair of cell types in each zone, we looked for an association of ligands in one cell type and all genes in the second cell type with the same directionality via the NicheNet algorithm. The background gene set for each cell type in each zone used in NicheNet was taken to be any gene expressed in at least 10% of target cells. We then only maintained ligands that achieved a NicheNet score > .1, and that were associated with receptors expressed in at least 30% of the target cell population. Lastly, for every ligand in a sending cell population, and associated target gene set in a target cell population, we queried all local neighborhoods of the target cells that belonged to the sending cell population (BFS on the Delaunay triangulation of cells with search depth of 2), and filtered for neighborhoods containing at least 3 cells, and where the total neighboring cells exceeded 500. With these sending cell neighborhood and target cell definitions, we ensured that ligand expression in the neighborhoods of the target cell population correlated with the target cells’ target gene signature scores. (i.e., Spearman correlation coefficient > 0 and p-value for non-correlation < .05).

We next defined ligand interactions with opposing directionality in infiltrated and non-infiltrated biopsies at baseline. Here, we first computed the differences in dot products with respect to the principal axis of each zone for each cell type between infiltrated and non-infiltrated biopsies at baseline. For every cell type in every zone at baseline, we took positive differences greater than the 95^th^ percentile of dot product differences to indicate genes that move more towards the principal axis of a zone in infiltrated than in non-infiltrated biopsies. Conversely, negative differences less than the 5^th^ percentile of dot product differences were considered genes that move more towards the principal axis of a zone in non-infiltrated biopsies. We further filtered gene sets based on genes expressed in at least 15% of each cell population in the CosMx data and in at least 2.5% of the cells’ cell type population in the scRNA-seq data. A cell population was only considered to be a target cell population if it contained more than ten genes in its signature. The same ligand filtering conducted in the ligand gradient analysis in each zone at baseline was used here.

Finally, we defined ligand interactions with unique directionality in each treatment compared to the previous treatment, in both infiltrated and non-infiltrated biopsies. For each cell type in each zone and in biopsies of each infiltration status, gene dot product differences were calculated between Base and PD1, PD1 and RTPD1. In B and T cell zones, for every cell type and treatment (in each comparison), genes with a significant gradient signature in one infiltration status and no significant signature or the opposite signature in the other were used as the gradient gene set. For malignant cell and intermediate cell zones, gradient genes were defined as those enriched in every cell type and treatment per comparison in non-infiltrated biopsies. The same target gene set and ligand filtering used previously in comparing infiltrated to non-infiltrated biopsies were used here.

### Secreted ligand-receptor analysis in cDCs describing shift in gene expression from baseline to PD1 in the MCZ

In non-infiltrated biopsies there is a relatively large population of cDCs which infiltrates the malignant zones after PD1. Any gene expression brought into these zones from cDCs would not show up in our earlier cell type specific differential expression analysis since cDCs were absent from the malignant cell zones at baseline. To study this we first performed differential expression on these malignant zones between baseline and PD1 for all cells, regardless of type, using the same cuts as described previously. We then performed the same isolation forest analysis on these genes to identify anomalous cell types with high expression. Secreted and membrane ligands were identified from the list of genes expressed by cDCs for analysis.

For secreted ligands shown to be brought to the malignant cell zone by cDCs… We used the same target gene set for cDCs in the malignant cell zone at PD1 compared to baseline in non-infiltrated biopsies that was defined previously in our localized ligand interaction analysis across infiltration and treatment statuses. Potential interacting ligands were found to be differentially expressed secreted ligands in all cells in the non-infiltrated malignant cell zones at PD1 compared to baseline. We honed in on a set of secreted ligands affecting the downstream target genes by applying the same differential expression thresholds and ligand prioritization used in the localized ligand interaction analysis. Sending populations were defined by the same anomaly detection protocol.

We then asked if there are ligands and receptors affecting the expression of CXCL9 in cDCs, the ligand found to be enriched in cDCs and affecting the genes enriched in T cells in the MCZ at PD1. To do so, we first looked for genes differentially expressed across all cells in the MCZ at PD1 compared to baseline, where FDR < .05, log2 fold change > .2 and were expressed at a rate of at least .15 in either group. Then, we filtered for cytokines and chemokines that were both differentially expressed, and associated with at least one receptor that was expressed in at least 30% of cDCs in the MCZ at PD1. We next identified which cell types were the most anomalous expressors of each chemokine or cytokine ligand. Additionally, we looked for ligands likely upstream of CXCL9. That is, we found ligands where CXCL9 was found in the genes corresponding with the top 5% of weights in the NicheNet database. Again, we filtered for those ligands that were differentially expressed across all cells in the MCZ at PD1, and expressed at least one receptor present in the cDCs in this zone and time point. Lastly, we identified the likely senders of these CXCL9-affecting ligands.

